# Telomouse – a mouse model with human-length telomeres generated by a single amino acid change in RTEL1

**DOI:** 10.1101/2021.06.06.447246

**Authors:** Riham Smoom, Catherine Lee May, Vivian Ortiz, Mark Tigue, Hannah M. Kolev, Melissa Rowe, Yitzhak Reizel, Ashleigh Morgan, Nachshon Egyes, Dan Lichtental, Emmanuel Skordalakes, Klaus H. Kaestner, Yehuda Tzfati

**Affiliations:** Department of Genetics, The Silberman Institute of Life Sciences, Safra Campus, The Hebrew University of Jerusalem, Jerusalem, 91904, Israel; Department of Genetics and Institute for Diabetes, Obesity and Metabolism, Perelman School of Medicine, University of Pennsylvania, Philadelphia, PA 19104, USA; The Wistar Institute, Philadelphia, PA 19104, USA; Division of Gastroenterology and Hepatology, Department of Medicine, Hospital of the University of Pennsylvania, Philadelphia, PA 19104, USA

**Author notes:** Correspondence (Y.T.) and (K.H.K.).

## Abstract

Telomeres, the ends of eukaryotic chromosomes, protect genome integrity and enable cell proliferation. Maintaining optimal telomere length in the germline and throughout life limits the risk of cancer and enables healthy aging. Telomeres in the house mouse, *Mus musculus*, are about five times longer than human telomeres, limiting the use of this common laboratory animal for studying the contribution of telomere biology to aging and cancer. We identified a key amino acid variation in the helicase RTEL1, naturally occurring in the short-telomere mouse species *M. spretus.* Introducing this variation into *M. musculus* is sufficient to reduce the telomere length set point in the germline and generate mice with human-length telomeres. While these mice are fertile and appear healthy, the regenerative capacity of their colonic epithelium is compromised. The engineered Telomouse reported here demonstrates a dominant role of RTEL1 in telomere length regulation and provides a unique model for aging and cancer.

## INTRODUCTION

Telomeres maintain genome stability and enable cell proliferation by repressing the DNA damage response at the ends of linear chromosomes^1^. Human telomeres in most somatic cells shorten with each cell division due to incomplete replication and degradation^2^. In the germline, however, telomeres are elongated by the enzyme telomerase to maintain them within a defined, species-specific range referred to as the “telomere length set point”^3^. The telomere length set point allows for sufficient, though not unlimited, cell divisions in the soma throughout life. Optimal age-appropriate telomere length is critical for maintaining genome stability and regulating cell proliferation and senescence, the major determinants of carcinogenesis and aging^4–8^. However, despite decades of research, it is not yet understood how telomere length homeostasis is achieved. The large difference (∼ five-fold) in telomere length between two closely related and inter-breedable species of mice, *Mus musculus* and *M. spretus*, was exploited previously to search for genes that determine species-specific telomere length^9–11^. Interestingly, none of the known telomere maintenance genes were identified in these screens. Rather, the difference in telomere length between the two species was attributed to a single locus encoding a DNA helicase, which was therefore termed ‘*Regulator of Telomere Length’* (*Rtel1*)^9, 11^. However, the specific difference between the *M. musculus* and *M. spretus* RTEL1 proteins that is responsible for the different telomere length set points has not been identified thus far^12^.

RTEL1 belongs to a small family of iron-sulfur helicases together with XPD, FANCJ and ChlR1, which were implicated in the human inherited diseases Xeroderma pigmentosum, Fanconi anemia, and Warsaw breakage syndrome, respectively ^13, 14^. Biallelic germline mutations in human *RTEL1* cause a fatal telomere biology disease named Hoyeraal-Hreidarsson syndrome (HHS)^15–19^. HHS is characterized by accelerated telomere shortening and diverse clinical symptoms including bone marrow failure, immunodeficiency, and developmental defects^20^. Other HHS-causing mutations were found in telomerase subunits, or in factors essential for telomerase biogenesis or recruitment to telomeres^21^. While RTEL1 was reported to have multiple roles in maintaining the stability of the genome, and particularly telomeres^22–27^, it was also suggested to regulate nuclear trafficking of ncRNA^28^, the localization of telomeric repeat-containing RNA (TERRA)^29^, and telomere elongation by telomerase^26, 30^.

One of the *RTEL1* missense mutations identified in HHS patients results in a single amino acid change from the highly conserved methionine 492 to isoleucine (Figure 1a,b)^17^. Methionine 492 is conserved nearly uniformly across vertebrates, with only a few species (mostly cats) having a leucine at this position. *M. spretus* is the only vertebrate species having a more radical change at this site, namely to the positively charged amino acid lysine^17^. We hypothesized that lysine 492 in *M. spretus* might be responsible for the difference in telomere length between the two mouse species. Here we describe an engineered mouse that confirms this hypothesis and presents an invaluable model for the study of cancer and aging.

**Figure 1.**
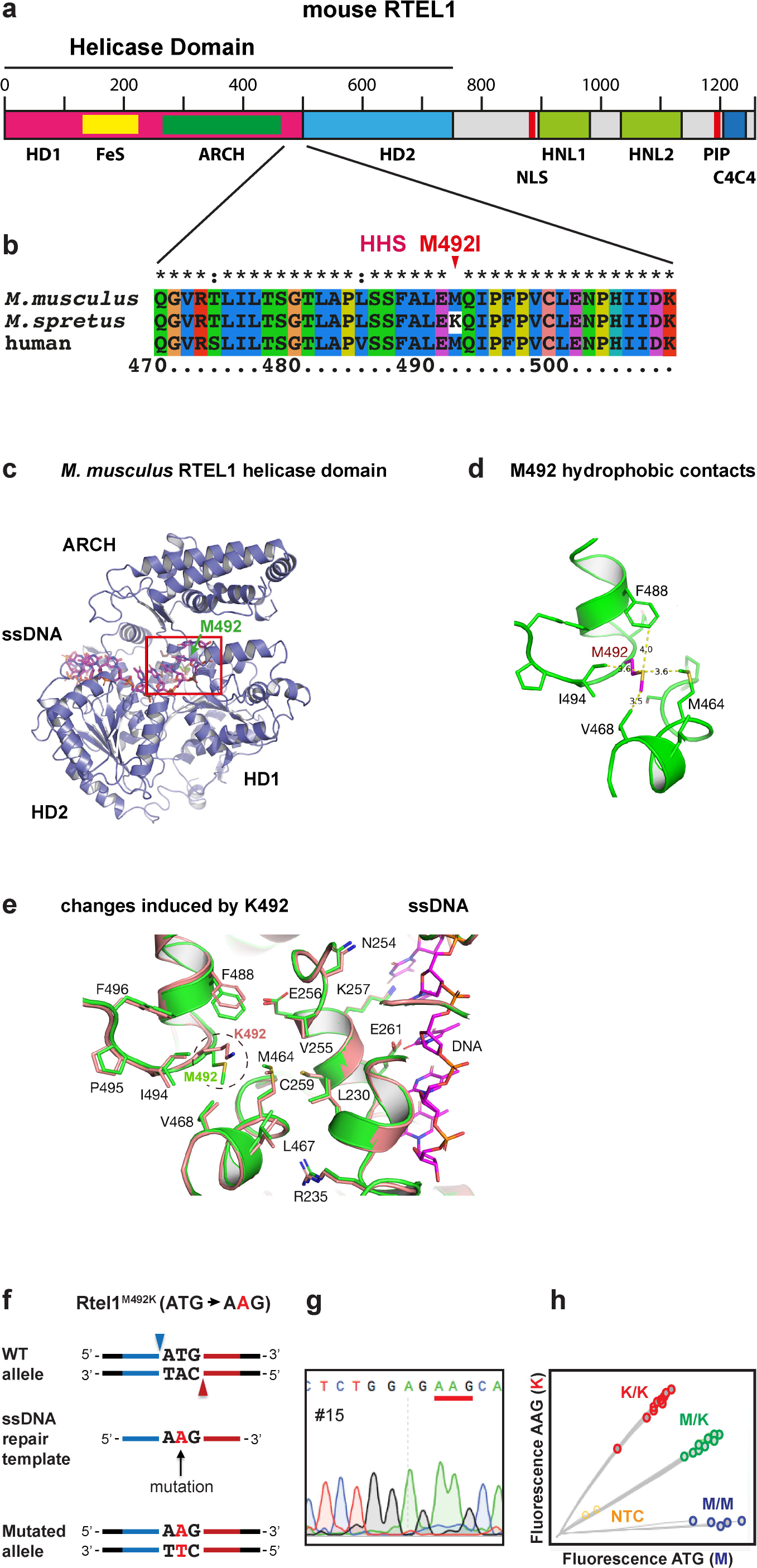
The RTEL1 helicase domain. (a) Map delineating the domains of RTEL1. The N-terminal part of the protein includes helicase domains 1 and 2 (HD1 and HD2), the iron-sulfur coordinating cluster (FeS) and the ARCH domain. The C-terminal part includes the nuclear localization signal (NLS), two harmonin N-like domains (HNL1 and HNL2), the PCNA interacting protein motif (PIP), and a C4C4 ring-finger domain. (b) Protein sequence alignment showing position 492 with a conserved methionine in *M. musculus* and human (mutated to isoleucine in HHS), and a lysine in *M. spretus.* (c) Predicted structure model of the *M. musculus* RTEL1 helicase domain, formed by residues 3-777 of mRTEL1 in a complex with single-stranded DNA. (d) Enlargement of the helicase domain in the red box in (c), showing the contacts of M492 with several hydrophobic residues and distances in Å. (e) Replacing M492 with the larger and charged lysine and overlaying the energy-minimized WT (green) and mutant (light brown) structures reveals changes in the position of nearby residues, which may affect the interaction with the adjacent single-stranded DNA. (f) Derivation of an *Rtel1*^M492K^ allele using the CRISPR-Cas9 nickase editing strategy. Illustration showing the WT allele, the two positions targeted by gRNAs (blue and red arrowheads), the ssDNA repair template with the mutation, left homology arm (blue), right homology arm (red) and the resulting targeted allele. (g) Sanger sequencing of founder #15 confirming the replacement of the “T” with an “A” at the codon encoding methionine 492, changing it to a lysine (underlined). (h) Genotyping of animals using a custom Taqman SNP Genotyping Assay. Fluorescence intensities of AAG (K) and ATG (M) are plotted for WT (M/M; blue), heterozygotes (M/K; green) and homozygotes (K/K; red) samples, and no-template control (NTC; yellow).

## RESULTS

### Methionine 492 is located at a critical position within the helicase core domain of RTEL1

Methionine 492 of RTEL1 is conserved in vertebrates but found to be altered in human HHS patients and in the short-telomere mouse species *M. spretus*^17^. To understand the role of this residue in RTEL1 function, we generated a structural model of the murine RTEL1 helicase domain using the Protein Homology/analogY Recognition Engine V 2.0 (Phyre 2)^31^ and then modeled the complex with single-stranded DNA based on the X-ray crystal structure of the bacterial XPD family helicase DinG in complex with the DNA^32^. This model revealed that methionine 492 (M492) is buried within the core of the helicase domain in proximity to the single-stranded DNA binding site of the protein, making extensive hydrophobic contacts with the hydrophobic side chains of residues M464, V468, F488, and I494 (Figure 1c,d)^32^. Replacing methionine with the larger and positively charged lysine within this hydrophobic patch is predicted to perturb the local structure (Figure 1e), which in turn may affect the helicase DNA binding, translocation and unwinding activities of the protein.

### A house mouse with short telomeres

To test if the lysine residue at position 492 of *M. spretus* is responsible for its shorter telomeres compared to the house mouse *M. musculus*, we generated a *M. musculus* strain in which methionine 492 was changed to a lysine (M492K) using CRISPR-Cas9 assisted genome editing in zygotes. We employed the Cas9 nickase (D10A Cas9 mutant) enzyme and two guide RNAs to generate nicks on both strands of the *Rtel1* gene to facilitate gene conversion using a repair template (Figure 1f). Successful gene targeting in the resulting offspring was confirmed by DNA sequencing and by allele-specific Taqman probe-based PCR (Figure 1g,h). This founder mouse was backcrossed to wild-type (WT) C57BL/6 mice to generate F0 mice heterozygous for the *Rtel1*^M492K^ allele, termed *Rtel1*^M/K^. F0 *Rtel1*^M/K^ mice were intercrossed to generate F1 and F2 *Rtel1*^M/K^. Homozygosity for the *Rtel1* mutation (*Rtel1*^K/K^) was established in the third generation (F3). The homozygous *Rtel1*^K/K^ mice were then bred *inter se* for thirteen additional generations up to F16 to date and were analyzed for telomere length and function. We termed the *Rtel1*^K/K^ mouse model ‘Telomouse’.

To determine the effect of the *Rtel1*^M492K^ mutation on telomere length, we employed two experimental paradigms. First, we established embryonic fibroblast cultures from *Rtel1*^M/K^ and *Rtel1*^K/K^ F3 embryos generated by intercrossing F2 *Rtel1*^M/K^ mice (Figure S1). As a control, we employed WT (*Rtel1*^M/M^) embryonic fibroblasts generated from intercrossing WT mice which did not have the *Rtel1*^M492K^ mutation in their pedigree, to avoid trans-generational inheritance of short telomeres from the heterozygous mice to the WT offspring, as occurred in the fibroblasts derived from the progeny of F2 *Rtel1*^M/K^ mice (M/M** in Figure S2a). Mouse embryonic fibroblast (MEF) cultures were established by serial passaging and selection for spontaneous immortalization and followed in culture over 250 population doublings (PD; Figure S1). During growth, the cultures passed one or two transition points in which the MEFs increased their growth rates and changed their cellular morphology, presumably by random inactivation of a tumor suppressor gene or genes such as p16 (Figure S1 and^33^). During the final growth phase, *Rtel1*^M/K^, *Rtel1*^K/K^ and *Rtel1*^M/M^ MEFs grew at the same average rate (about one PD/day). We determined the average telomere lengths and the telomeric 3’-overhang length for these three cultures at multiple PD levels by the in-gel hybridization method as follows. We digested high molecular weight genomic DNA samples with the restriction endonuclease *HinfI*, separated the restriction fragments by pulse-field gel electrophoresis (PFGE) and dried the gel. First we hybridized the native DNA in the gel to a telomeric C-rich probe to estimate the average length of the G-rich single-stranded telomeric overhang (see below). Then, the DNA was denatured and hybridized again to the same probe to detect both single- and double-stranded telomeric repeat tracts and estimate the length of the telomeric restriction fragments (TRF; representative gel images are shown in Figures 2a, S2 and S3). The mean TRF length (MTL) of each sample was repeatedly measured in different gels to increase the measurement precision (Summarized in Table S1 and Figure 2b). The average MTL in WT (*Rtel1*^M/M^) MEFs was 46.9 kb at PD 10. It shortened to about 40 kb during the initial growth phase (Figures S1 and 2b), presumably due to the lower telomerase activity present in MEFs prior to immortalization^36^. Upon immortalization at about PD 70 (indicated by arrows in Figure 2a,b), telomere length stabilized, reaching an average MTL of 37.8 kb at PD 250. In contrast, the mutant *Rtel1*^M/K^ and *Rtel1*^K/K^ MEFs displayed shorter telomeres at PD 10 (average MTL 34.2 and 30.1 kb, respectively), which continued to shorten gradually even beyond the early growth phase at an average rate of about 50 bp per PD, to 21.6 and 14.7 kb at PD 250, respectively (Figure 2a,b). Altogether, the average MTL in the *Rtel1*^K/K^ MEFs shortened by 60% as compared to the WT TRF length after 250 PD, and reached a length comparable to that of human fibroblasts immortalized by ectopic expression of telomerase^34^.

**Figure 2.**
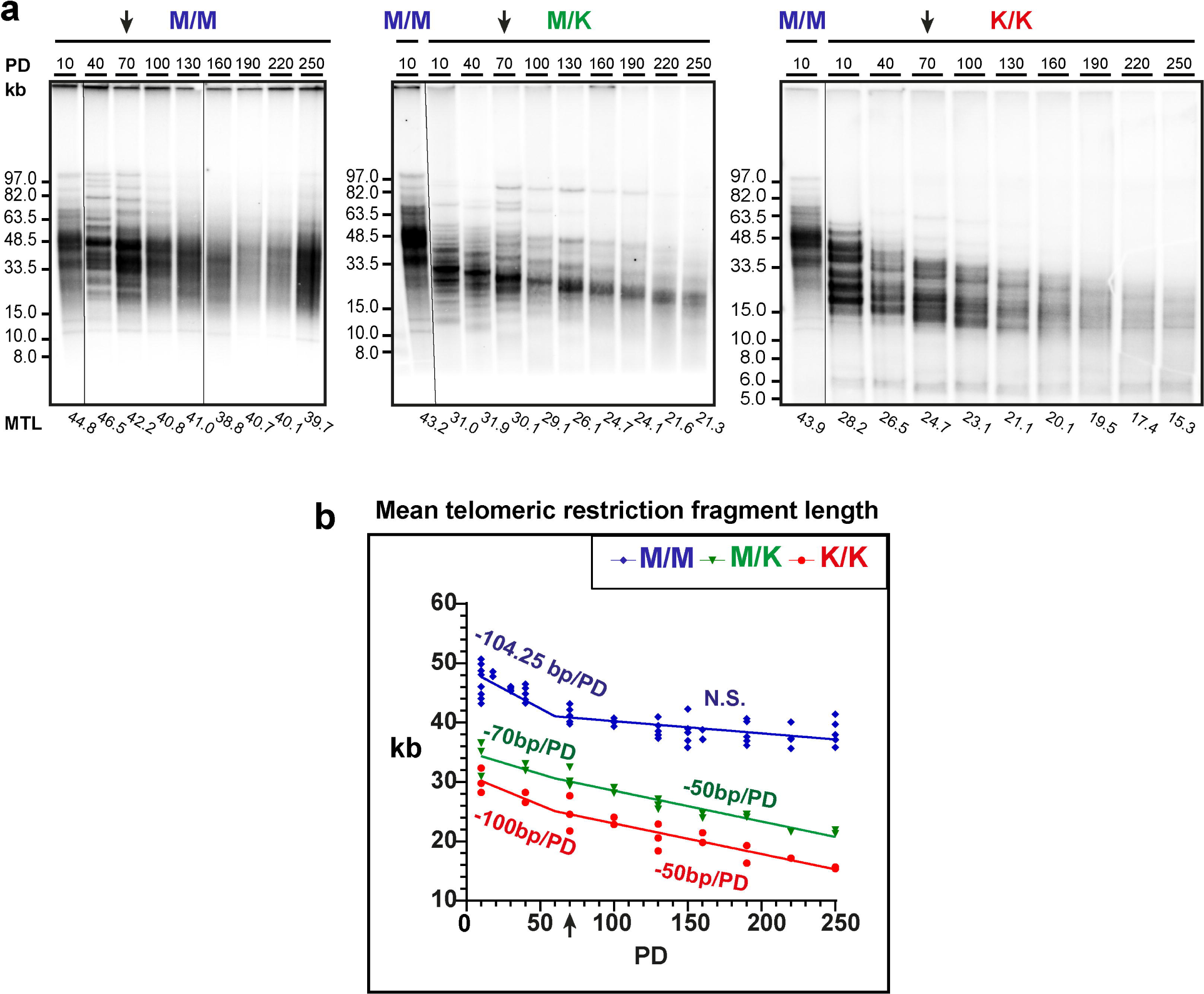
Telomeres of MEFs carrying the *Rtel1*^M492K^ mutation shorten progressively with cell division in culture. (a) Genomic DNA samples prepared from WT (M/M) or mutant MEF cultures homozygous (K/K) or heterozygous (M/K) for *Rtel1*^M492K^, were analyzed by PFGE and in-gel hybridization to the denatured DNA. The un-spliced image of the gels are shown in Figure S2a,c,e. Mean telomere length (indicated below the lanes) was quantified using *TeloTool* ^51^. (b) Mean TRF length for each of the sampled PD were repeatedly measured in additional gels (summarized in Table S1) and plotted. N.S. indicates no significant difference from a horizontal line (slope 0). The complete change of cell morphology reflecting cellular immortalization (see Figure S1c) is indicated by arrows on the gel images (a) and graph (b).

RTEL1 dysfunction was proposed previously to accelerate telomere shortening either by telomere deletion or by reducing telomerase action at telomeres^25, 26, 30^. To investigate the cause for telomere shortening in *Rtel1^K/K^* MEFs, we examined the distinct TRF banding pattern appearing in the *Rtel1^M/K^* and *Rtel1^K/K^* MEFs (Figures 2a, S2 and S3). Such banding pattern was observed previously in telomerase null MEFs^35^, as well as in mRtel1-deficient mouse embryonic stem cells^26^. These types of bands were suggested to represent homogenous telomere lengths of individual chromosomes that became clonally fixed within the cell population because they did not undergo significant elongation despite the presence of telomerase^26^. Furthermore, these TRF bands showed gradual telomere shortening over time in culture, indicating the inability of telomerase to compensate for the telomere shortening that occurs with each round of cell division^26^. Based on this observation, Uringa and colleagues hypothesized that mRtel1 was required for telomere elongation by telomerase. Similarly, *Rtel1*^M/K^ and *Rtel1*^K/K^ MEFs maintained several such sharp bands, which gradually shortened with continued passaging (Figures S2 and S3), indicating that telomerase activity was insufficient to maintain stable telomere length in these cells. Calculating the rates of shortening of individual bands reveal faster shortening of up to 170 bp per PD during the initial growth phase, and slower shortening of up to 66 bp per PD during the later phase in both heterozygous and homozygous mutant MEFs (Figures S2b,d and S3d). WT MEFs also displayed sharp bands during the initial growth phase (Figures 2a, S2e,f and S3c,d). However, these bands did not appear to shorten but rather disappeared upon immortalization, presumably due to the increased telomerase activity associated with immortalization of MEFs^36^. These findings indicate that the shortening of telomeres in *Rtel1*^K/K^ and *Rtel1*^M/K^ MEFs was not due to random telomere deletion, which would have caused these bands to disappear, but rather the result of reduced telomerase action at telomere ends.

It was proposed previously that the actual length of the telomeric repeat tracts in mouse telomeres is much shorter than the length of the TRF observed by gel electrophoresis due to low sequence complexity and limited representation of restriction sites in the subtelomeric regions^35, 37^. However, the exact length of the subtelomere included in the TRF is still unknown and expected to vary between chromosome ends and different restriction enzymes used to digest the genomic DNA to be analyzed. To try to estimate the average length of the subtelomeric regions within the TRF we examined the correlation between the hybridization signal and mean TRF length. Since equal amounts of genomic DNA were loaded for each sample on the gels, we expected the denatured hybridization signal within each gel to correlate with telomere length, as longer telomeres bind more molecules of the oligonucleotide probe and thus have stronger signals. Plotting mean TRF length as a function of the telomeric hybridization signal indeed revealed a linear correlation (Figure S4). Extrapolation of the regression line to zero signal did not reach zero TRF length as expected if the TRF sequence were entirely telomeric, but rather suggested that the observed TRFs are longer than the telomeric repeat tracks by an average of 10-15 kb (Figure S4). This analysis confirmed that at least some chromosome ends have long subtelomeric sequences of low complexity that are underrepresented for *HinfI* restriction sites, contributing to the measured TRF length. Therefore, the actual lengths of the telomeric repeat tracts are significantly shorter than appears by PFGE, as proposed previously^35, 37^. TRFs with very short telomeres may not be detected by gel hybridization even if their subtelomeric portions are long because the telomeric hybridization signal is too low. In addition, very short TRFs may not be detected because they fall outside the range observed by PFGE.

To directly compare the actual length of the telomeric repeat tracts between mouse and human telomeres, we used Quantitative Fluorescence In Situ Hybridization (qFISH). We mixed together metaphase-arrested *Rtel1*^K/K^ MEFs at PD 250 with human telomerase-positive fibroblasts, spread them onto slides and hybridized them to telomeric (green) and centromeric (red) peptide nucleic acid (PNA) probes (Figures 3a and S5). *Rtel1*^M/M^ control MEFs prepared and hybridized under the same conditions are shown for comparison in Figure 3b. The mean TRF lengths as calculated by gel electrophoresis for the *Rtel1*^K/K^ MEFs and the human fibroblasts were 14.7 kb and 14 kb, respectively (Figures 2b and 3d). The centromeric probe enabled us to distinguish the human chromosomes, which are mostly metacentric, from the acrocentric mouse chromosomes. The telomeric signals at each chromosome end were quantified using the *Telometer* plugin (available at https://demarzolab.pathology.jhmi.edu/telometer/index.html) of *NIH ImageJ*^38^. The average telomeric signal intensity of the *Rtel1*^K/K^ MEFs at PD 250 was comparable or lower than that of the human telomeres (Figures 3a,c and S5a,d). In addition, the distribution of individual telomeric signals was significantly wider in the MEFs with a significant portion of telomeres below the detection limit (Figure 3c). Most of these telomeres displayed very weak hybridization signals, detectable only under prolonged exposure and increased gain, indicating that they were extremely short but not completely lost (Figure S5e). Moreover, regular (constant field) gel electrophoresis followed by in-gel hybridization revealed that a significant fraction of TRF in the *Rtel1*^K/K^ MEFs PD 250 was shorter than 5 kb (Figure 3d). In sum, these data are consistent with previous publications suggesting that long subtelomeric regions are included in the TRF due to low sequence complexity and that a significant distance exists between the telomeric repeat tracts and the most distal restriction site^35, 37^. The position of the restriction site likely varies between chromosome ends, thus contributing to the heterogeneity of TRF lengths observed by PFGE. As the telomeres of the *Rtel1*^K/K^ MEFs shorten, the error introduced by the variable subtelomeric portion in the TRF measured by PFGE becomes even more significant in determining the actual length of the telomeres. In addition, some TRF with short subtelomeric regions become so short that they are excluded from the size range detected by PFGE. These conclusions highlight the need for a more accurate method for measuring the actual length of the telomeric repeat tracts.

**Figure 3.**
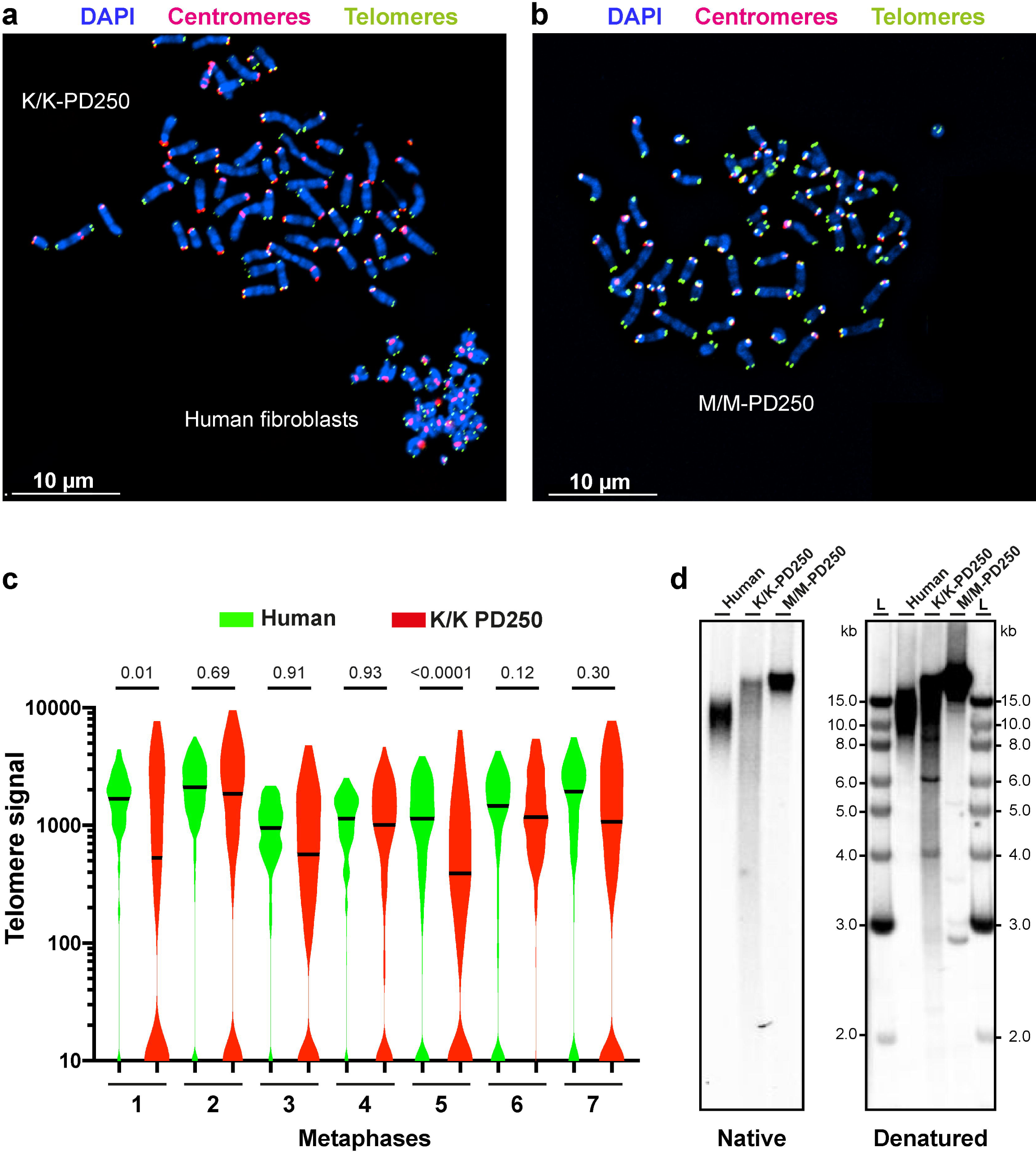
The *Rtel1*^K/K^ MEFs telomeres are comparable or shorter to those of human telomeres. (a) *Rtel1*^K/K^ MEFs PD 250 (with a mean TRF length of 14.7 kb measured by PFGE and in-gel hybridization) and telomerase-positive human primary fibroblasts (with a mean TRF length of 14 kb measured by in-gel hybridization) were arrested in metaphase, mixed, and spread on slides. The slides were hybridized to a green telomeric PNA probe and a red centromeric probe to distinguish the human mostly metacentric chromosomes from the mouse acrocentric chromosomes. (b) *Rtel1*^M/M^ MEFs PD 250 were processed as in (a), under the same conditions and microscope settings by a Nikon Eclipse Ti-E microscope equipped with a CoolSNAP Myo CCD camera.. (c) The telomeric signals were quantified using the *Telometer* plugin of *NIH ImageJ*^38^. Violin plots show the distribution of individual telomeric signal intensities in seven microscope fields containing pairs of human and mouse *Rtel1*^K/K^ metaphase cells. Additional pairs are shown in Figure S5. For the human samples between 116 and 324 chromosome ends were quantified per field. For the mouse samples, between 178 and 248 chromosome ends were analyzed per field. The higher number of telomeres per field than the expected 184 (human) and 160 (mouse) may result from more than one metaphase combined in the same field or aneuploidy. Horizontal lines indicate the median. Unpaired two-tailed t-test showed significant differences in the telomeric signals between the *Rtel1*^K/K^ MEFs PD 250 and human fibroblasts in pairs 1 and 5. The P values are indicated above each pair. Nested one way ANOVA for all seven pairs indicate significantly lower signal intensity in the MEFs (P = 0.003). (d) Genomic DNA was extracted from the same cultures shown in (a) and (b) and analyzed by regular (constant field) gel electrophoresis and in-gel hybridization to the native and denatured DNA.

### NanoTelSeq - a novel method for single telomere analysis by long-read Nanopore sequencing

It is well established that the shortest telomeres are the most biologically relevant as they may activate the DNA damage response and determine cellular function and fate. However, as apparent from the results detailed above, currently used methods based on measuring hybridization signals or telomeric restriction fragments fail to measure the shortest telomeres accurately. A method called TeSLA was designed to PCR-amplify and measure the shortest telomeres^39^. However, it is labor intensive, has significant limitations, and is practically impossible to use for analyzing the ultra-long telomeres of *M. musculus*. To address this major shortfall, we developed a method for single telomere analysis by long-read Nanopore sequencing and termed it *NanoTelSeq*. In this method we adapted a previously developed approach to ligate a specific telorette linker with a C-rich telomeric repeat, which base-pairs with the G-rich telomeric overhang and can ligate to the 5’ end of the telomere and enable PCR amplification of the telomere^40^. In our method, a double-stranded telorette is not used for PCR amplification but enables the specific attachment of the Nanopore sequencing adapter to the 5’ end of the telomere (Figure 4a). The sequencing adapter facilitates the sequencing of the telomeric C-rich strand directly from 5’ to 3’ without any PCR amplification or selection to avoid bias. The presence of the telorette linker at the 5’ end of the sequence obtained, which requires an intact telomeric 3’ overhang for ligation, and the presence of a subtelomeric sequence at the 3’ end of the sequence reads, enabled us to accurately determine the complete length of the telomeric repeat tracts of individual telomeres. Using *NanoTelSeq* we sequenced undigested high molecular weight genomic DNA samples prepared from the *Rtel1*^K/K^ and *Rtel1*^M/M^ MEFs at PD 250 as well as human telomerase-positive skin fibroblasts and obtained several hundreds of telomere reads each (Figure 4). We first employed a mixture of six telorettes with the six possible permutations of the six nt telomeric repeat at their 3’ end. As reported for human telomeres^40^, telorette 3 was the most frequently used (60 % of the *Rtel1*^M/M^ and 76 % of the *Rtel1*^K/K^ and the human telomeric reads), indicating that the 5’ strand of most telomeres ends in 3’-CCAATC-5’ (Figure 4b). This specificity can only be expected if the telorette adapter was attached to native telomere ends and not to randomly broken telomeres, thus ensuring that most telomere reads represent intact telomeres. Therefore, for all additional sequencing reactions we employed only telorette 3, which was attached to different barcodes. For long reads that begin with the telorette sequence on the 5’ end of the telomeric C-rich strand and terminate with a subtelomeric sequence on the 3’ end, we could therefore determine the length of the telomere repeat sequences at base resolution. Telomeric reads were identified in the sequencing output data, processed and presented graphically by a dedicated computer program termed *NanoTel* (examples for telomere reads are shown in Figure 4c; all telomeric reads are summarized in Table S2). The median read length (N50) was between 20 and 50 kb, with longer reads up to about 200 kb. Since the length of the mouse telomeres is within the same range of the read length, not all reads reached the subtelomere (referred to as ‘truncated telomeres’). We wished to include as many telomere reads as possible to increase accuracy, without introducing a significant error in the median telomere length calculations due to the inclusion of truncated telomere sequences. We hypothesized that inclusion of sequences with a read length longer than the median telomere length would not introduce an error in the median telomere length calculation. To test it we used the samples with the highest number of telomere reads to compare median telomere length values calculated for all telomere reads, for all reads longer than the median telomere length, and for all reads longer than the longest incomplete read. Presumably, the latter represents the most accurate calculation, if the sample size was large enough. As shown in Figure S6, excluding all reads shorter than the median telomere length (middle plot for each sample) results in a significantly longer median telomere length, as compared with the median calculated for all telomeric reads (left), and very close to the value calculated for the reads longer than the longest incomplete (right). Only for MEFs M/M PD 250, the median telomere length calculated for the reads longer than the longest incomplete (right) was smaller than the value calculated for all reads longer than the middle telomere length (middle), presumably because of a too small sample size. Using the telomeric reads longer than the median, we found that the median length of the telomeric repeat tracts in *Rtel1*^K/K^ MEFs PD 250 was 5.59 kb, which is about 5 times shorter than the telomeres of WT MEFs (median length of 27.3 kb), and also shorter than the telomeres of human telomerase-positive fibroblasts (median length of 9.47 kb; Figure 4d). Furthermore, 19.5% of the telomeres of the *Rtel1*^K/K^ MEFs PD 250 were shorter than 3 kb, and 6.3% were below 2 kb, while only one of 602 telomere reads of the *Rtel1^M/M^* MEFs was shorter than 3 kb and none were shorter than 2 kb (Figure 4d). Nanopore sequencing thus enables the determination of telomere length at base resolution and, most importantly, the identification and analysis of abundant very short telomeres, which are not represented in the standard PFGE TRF or qFISH analyses but are nevertheless of utmost importance in determining cell function and fate.

**Figure 4.**
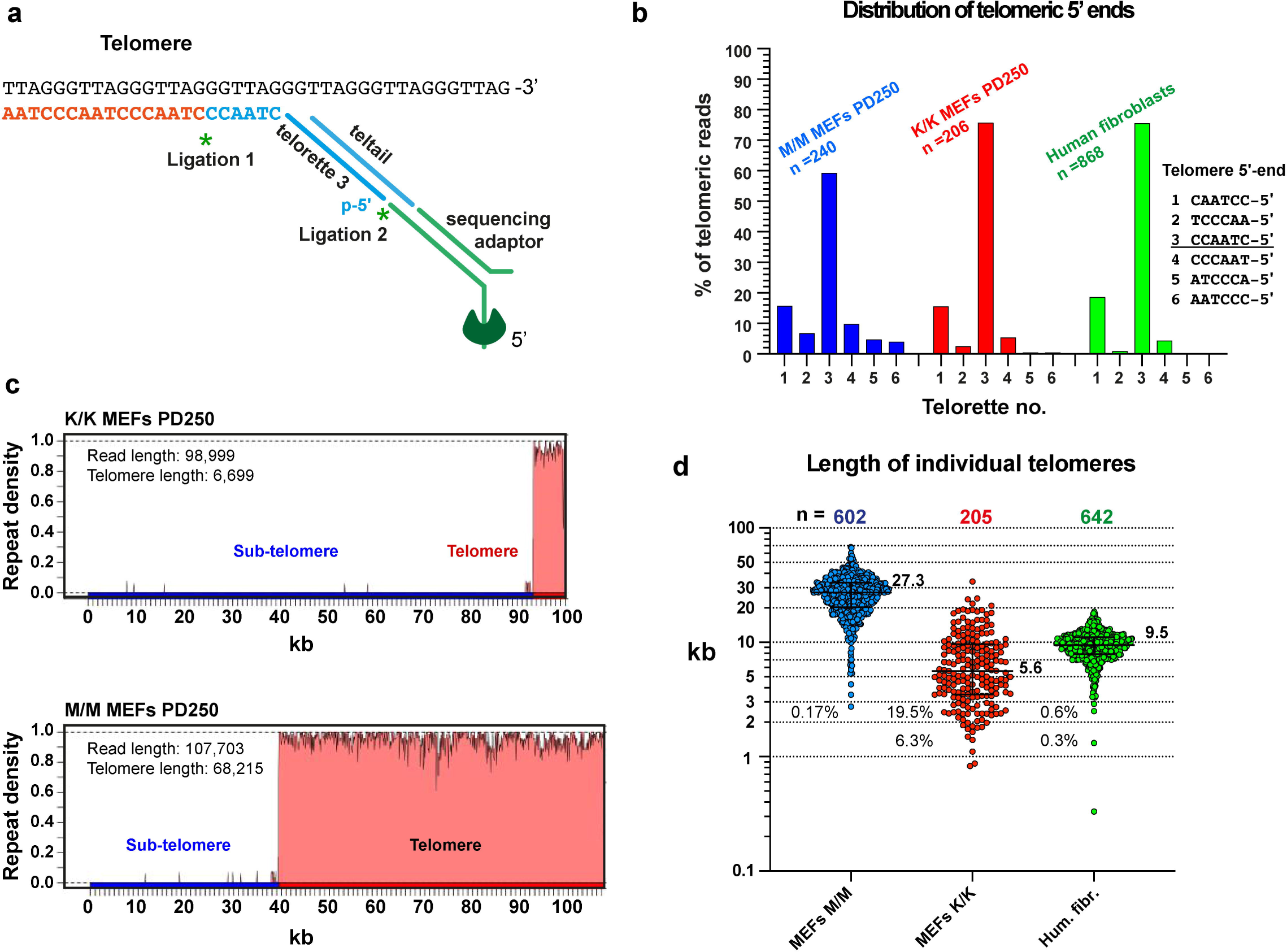
Long-read Nanopore sequencing reveals length heterogeneity and short telomeres in the *Rtel1*^K/K^ MEFs. (a) A scheme showing a telorette oligonucleotide complementary to the G-rich telomeric overhang ligated to the 5’ end of the telomere, followed by the ligation of the Nanopore sequencing adapter. (b) High molecular weight genomic DNA samples prepared from *Rtel1*^K/K^ and *Rtel1*^M/M^ MEFs PD 250 and human telomerase positive fibroblasts were ligated to a mixture of equal amounts of six telorette oligonucleotides, each ending with a different permutation of the C-rich 6 nucleotide telomeric repeat, and then ligated to the Nanopore sequencing adapter and sequenced each separately. The bar graph shows the percentage of the telomeric reads ligated with each telorette. (c) The reads were identified and processed computationally, and presented in graphs of telomeric repeat density (from 0 to 1, within a moving window of 50 nucleotides). Two representative examples are shown and the other reads are summarized in Table S2. Indicated are the precise telomere length and read length. (d) Scatter plots show the length of individual telomeres of the M/M and K/K MEF cultures at PD 250 and telomerase-positive human fibroblasts for comparison (the same cultures are shown in Fig3), with median and quartiles indicated by horizontal lines. Median values in kb are indicated to the right of each scatter plot and the percentage of telomeres below 3 kb and 2 kb to the left of the plots.

### Functional short telomeres in *Rtel1^K/K^* MEFs

The unexpected finding of telomeres below 3 kb (19.5%) and even 2 kb (6.3%) in the *Rtel1*^K/K^ MEFs at PD 250 raised the question whether these short telomeres are functional. The main role of telomeres is to suppress the DNA damage response (DDR) at chromosome ends and to prevent deleterious fusions of chromosomes and cell cycle arrest^1^. The presence of DDR foci at telomeres, termed telomere dysfunction-induced foci (TIF), is a hallmark of telomere failure. To examine if the telomeres of the *Rtel1*^K/K^ MEFs retained their protective properties, we studied the presence of TIF using antibodies against the DDR marker γH2AX and the telomere protein TRF1 (Figure 5a). The overall levels of DDR were high in all MEF cultures, as reported previously for MEFs growing under atmospheric oxygen levels^41^. However, there was no significant difference in the number of genome-wide DDR or TIF foci between WT (M/M) and mutant (K/K) MEFs (Figure 5a). To identify more accurately all the DDR foci associated with chromosome ends (even those without a detectable telomeric signal) we combined FISH and Immunofluorescence (IF) on metaphase chromosomes (Figure 5b). This analysis again revealed no significant difference in TIF formation between the genotypes. Furthermore, there was no increase in DDR even at those chromosome ends which lacked a detectable telomeric hybridization signal (‘signal-free ends’; Figure 5b), indicating that even the very short *Rtel1*^K/K^ telomeres that are below the limit of detection by FISH remained functional and able to suppress the DDR.

**Figure 5.**
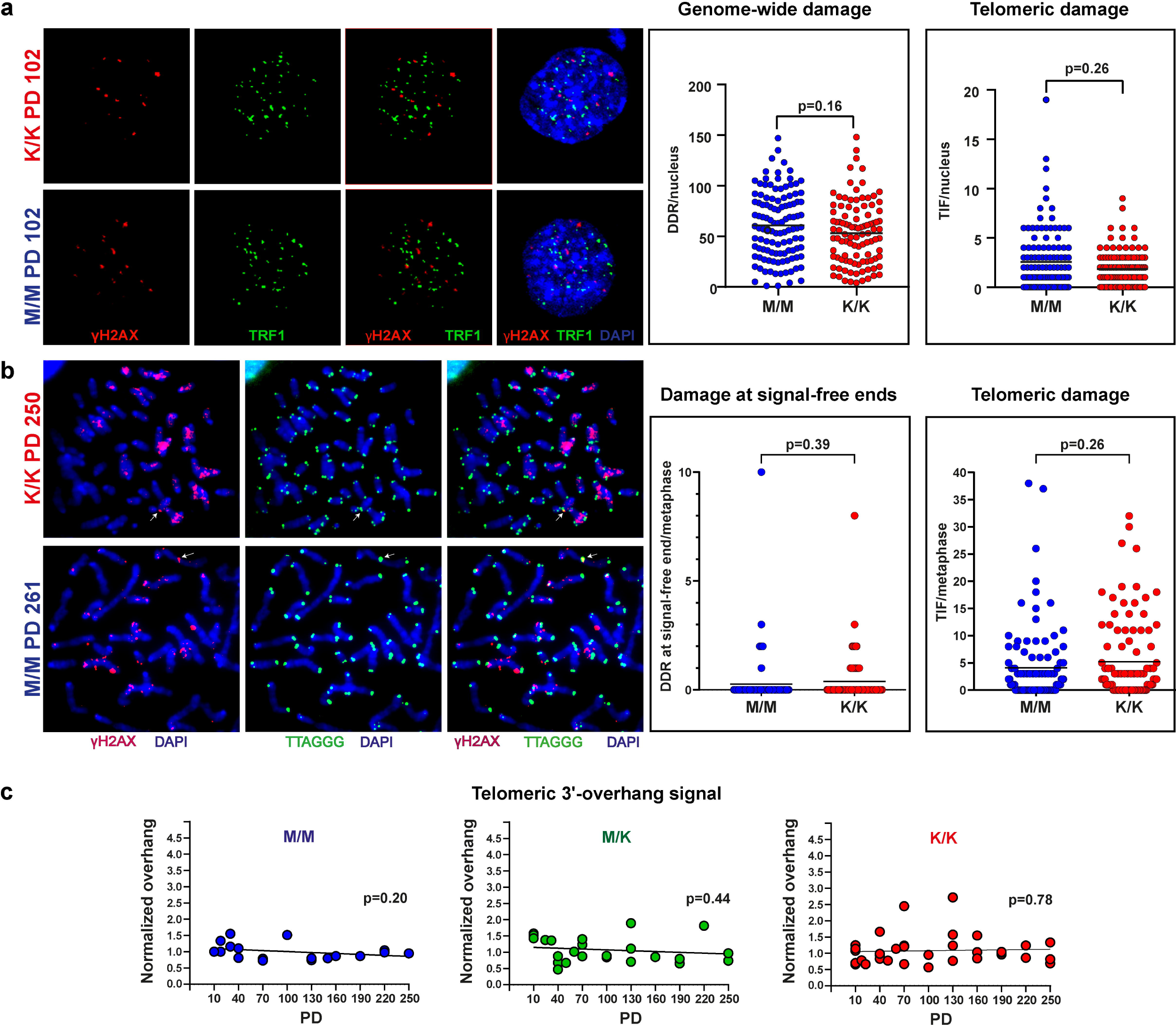
The short telomeres of *Rtel1*^K/K^ MEFs retain their protective function. (a) The formation of DDR foci and their localization to telomeres (defined as TIF) in interphase nuclei were evaluated by IF with antibodies detecting the DDR marker γH2AX (red) and the telomere protein TRF1 (green). Scatter plots show the number of DDR foci and TIF per nucleus. A total of 127 and 111 nuclei were counted for M/M and K/K MEFs (PD 102), respectively, in three independent experiments. The mean values are indicated by black lines, and P values were calculated by a nested t-test. (b) Metaphase TIFs were analyzed by FISH-IF using a γH2AX (red) antibody and a telomeric C-probe (green). A total of 100 metaphases were counted for each M/M (PD 261) and K/K (PD 250) MEF population in two independent experiments. The mean values are indicated by black lines, and P values were calculated by a nested t-test. (c) The normalized native overhang signal for 77 samples was measured and plotted as described in Figure S7. The slopes for K/K, M/K, and M/M were analyzed by simple linear regression and P values were calculated for the deviation from a horizontal line.

Shortened telomeric overhangs were found to be associated with telomere dysfunction in HHS patients and heterozygous carriers of the pathogenic M492I mutation in *RTEL1*^17, 30, 42^. To test whether the *Rtel1*^M492K^ mutation affects telomeric overhang maintenance in MEFs, we quantified the native in-gel hybridization signal and normalized it to a WT sample (M/M PD 10) as a measure of the relative average length of the telomeric overhang. The native signal remained constant and comparable to that of WT fibroblasts, indicating no significant impact of the Rtel1M492K mutation on the overhang maintenance (Figures 5c and S7).

RTEL1 dysfunction in patient cells and mouse *Rtel1* null cells was reported to induce telomere aberrations observed by telomere FISH on metaphase chromosomes, such as telomere loss, telomere fragility, interstitial telomere insertion (ITS) or telomere fusion^17, 25, 30^. To examine the frequency of telomere aberrations, we performed metaphase FISH on MEF cultures at PD 100 and 250 (Figure 6a,b,c). The main phenotypes observed were telomere loss and Robertsonian fusions, which increased from PD 100 to 250 in association with the shortening of the telomeres (Figure 2). Interestingly, most undetected telomeres by PFGE or qFISH (Figures 2a and 3a,c) actually had weak telomeric signal observed under increased exposure (see long arrows in Figure S5e), in line with the short telomeres measured by *NanoTelSeq* (Figure 4d). The few telomeres without any detectable signal, even under increased FISH exposure, underwent chromosome fusion (short arrows in Figure S5e). Presumably, Robertsonian fusions between the ends of the short p arms (but not the long q arms) are tolerated by the cells because of the proximity of the acrocentric centromeres to the p end, and can accumulate, unlike other deleterious fusions (Figure 6b,c). Other aberrations did not accumulate significantly more in the K/K mutant than in the WT control cells (Figure 6b,c). Finally, telomere sister chromatid exchange (T-SCE), indicating elevated telomeric recombination, was reported in RTEL1-deficient patient cells^16^. We examined the frequency of T-SCE events in the mutant cells by chromatid-orientation (co)-FISH, differentially labeling leading and lagging telomeres with green and red strand specific PNA probes. T-SCE events appear as colocalization of the two probes at the same telomere end (Figure 6d). There was no increase in T-SCE in the K/K mutant compared with the WT MEFs, indicating no increase in homologous recombination at the telomeres of the mutant K/K MEFs. Altogether, although the K/K MEFs displayed dramatically shortened telomeres, these telomeres largely maintained normal telomeric overhangs, which are essential for telomere end structure and function, and retained the ability to suppress the DDR and homologous recombination. Only Robertsonian fusions significantly accumulated in the K/K MEFs due to increased occasional telomere loss.

**Figure 6.**
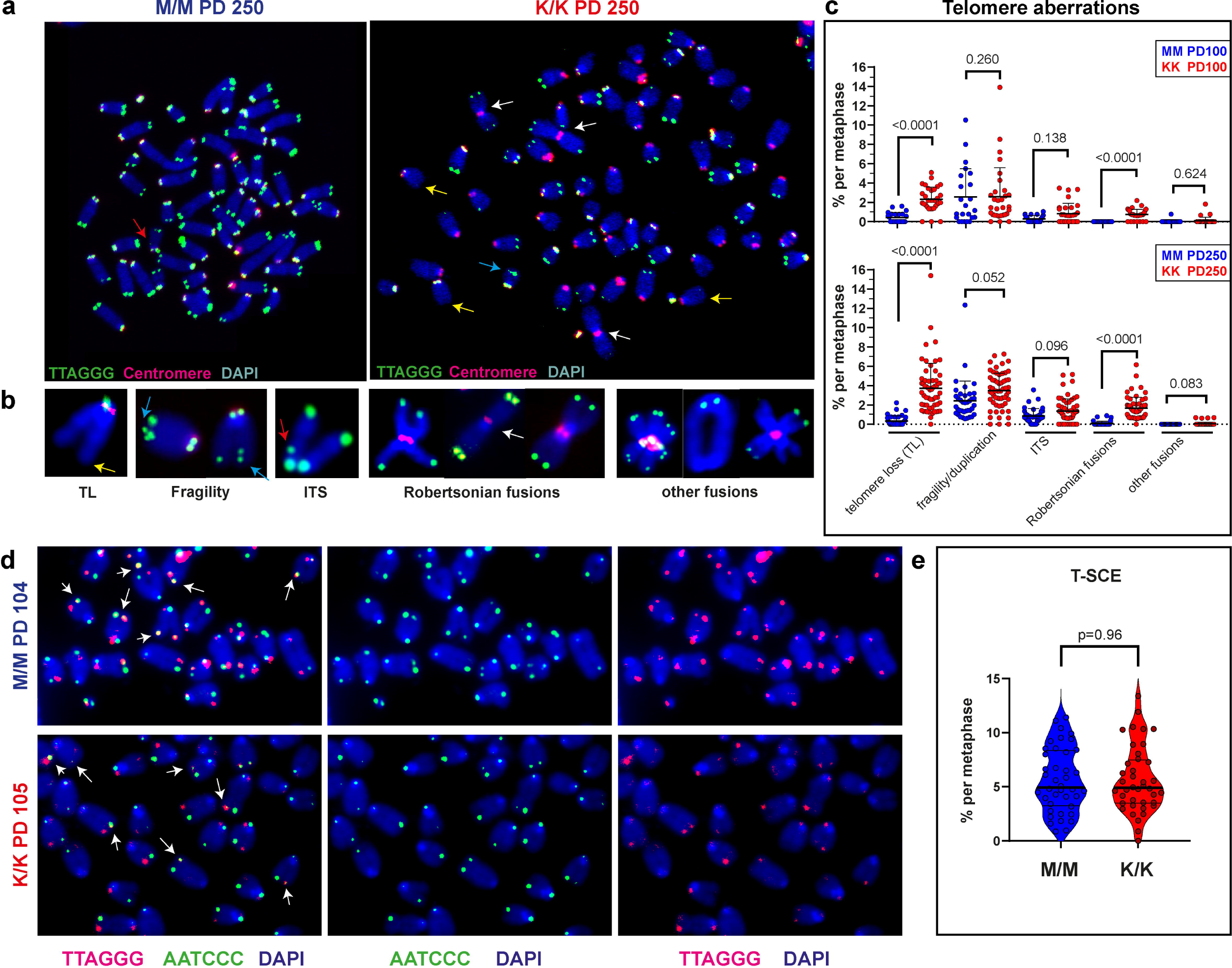
The chromosomes of *Rtel1*^K/K^ MEFs display increased telomere loss and Robertsonian fusions. (a) Representative images for metaphase chromosomes of M/M and K/K MEFs at PD 250 hybridized with telomeric (green) and centromeric (red) probes. Arrows indicate chromosomal aberrations. (b) Aberration types are shown enlarged: telomere loss (TL), telomere fragility, interstitial telomere insertions (ITS), Robertsonian fusions and other fusions. (c) Scatter plots show the percentage of each aberration type per metaphase (normalized to the number of chromosomes ends detected) at PD 100 (top) and PD 250 (bottom). Horizontal lines show mean and standard deviation. Two independent experiments for each M/M PD 100 and K/K PD 100 (n = 20 metaphases each experiment), and three independent experiments for each M/M PD 250 and K/K PD 250 (n = 35 metaphases each experiment) were performed. P-values were calculated by two-tailed t-test. (d) Chromatid-Orientation (CO)-FISH was performed on K/K and M/M MEFs at PD 105 and PD 104 respectively, labeling the leading and lagging telomeres with green and red using strand-specific PNA probes. Two experiments were done each for M/M and K/K (n = 20 metaphases each experiment). Telomere sister chromatid exchange (T-SCE) events (indicated by arrows) were counted and presented as a percentage of all telomere ends available for analysis in each metaphase. Horizontal lines show mean and standard deviation. P-value was calculated by two-tailed unpaired t-test.

### Short telomeres in Telomice

Telomeres of the *Rtel1*^K/K^ MEFs shortened over time in culture, as described above. However, telomere shortening in cultured fibroblasts does not necessarily imply reduced telomere length in the germline or in somatic tissues of mice. Therefore, we next analyzed telomere length in blood and tail DNA samples taken from *Rtel1*^K/K^ Telomice over 13 generations (F4 to F16). In the blood, the mean TRF length shortened to an average length of 16.7 kb in F16 Telomice, as compared to 39.7 kb in the WT samples (Table S3 and Figure 7a,c). In tail samples, mean TRF length shortened to an average of 19.2 kb in F16 Telomice as compared to 42.6 kb in the WT samples (Table S3 and Figure 7b,d). Importantly, when comparing the average TRF lengths of only the last two generations, F15 and F16, no further shortening was observed, neither in blood nor in tail samples, suggesting that telomere length had stabilized at a new telomere length set point in F16 mice (Figures 7c,d and S8). Overall, the TRF lengths shortened in the blood and tail at about 1 kb per generation. In comparison, telomeres of *M. musculus* deleted for telomerase RNA (*mTR^-/-^*) shorten at a much faster rate of 4.8 ± 2.4 kb per generation^35^. These results indicate that telomerase activity was not completely eliminated in Telomice but only down-regulated to reach a shorter telomere length set point. Consistent with the observation in MEFs, no reduction was observed in the native gel hybridization signal corresponding to the single-stranded G-rich telomeric sequences (Figures S9 and S10, for tail and blood samples, respectively), indicating that Telomice maintained an intact telomere end structure. Importantly, when we analyzed telomere repeat length by *NanoTelSeq*, we found Telomice (F15) to have a median repeat lengths of 6.4 kb in the blood and 6.1 kb in tail DNA, which is comparable to that measured in a human blood DNA sample (6.5 kb; Figure 7e). While only one telomere of the WT mouse blood sample was shorter than 3 kb, 4.6 % of the telomeres in the F15 Telomouse blood were shorter than 3 kb (as compared to 3% observed in human blood). Furthermore, 2.3 % and 3.4% of the telomeres of the Telomouse F15 blood and tail (respectively) were below 2 kb (Figure 7e). These observations suggest that short telomeres may play an important role in Telomice physiology but not in WT mice.

**Figure 7.**
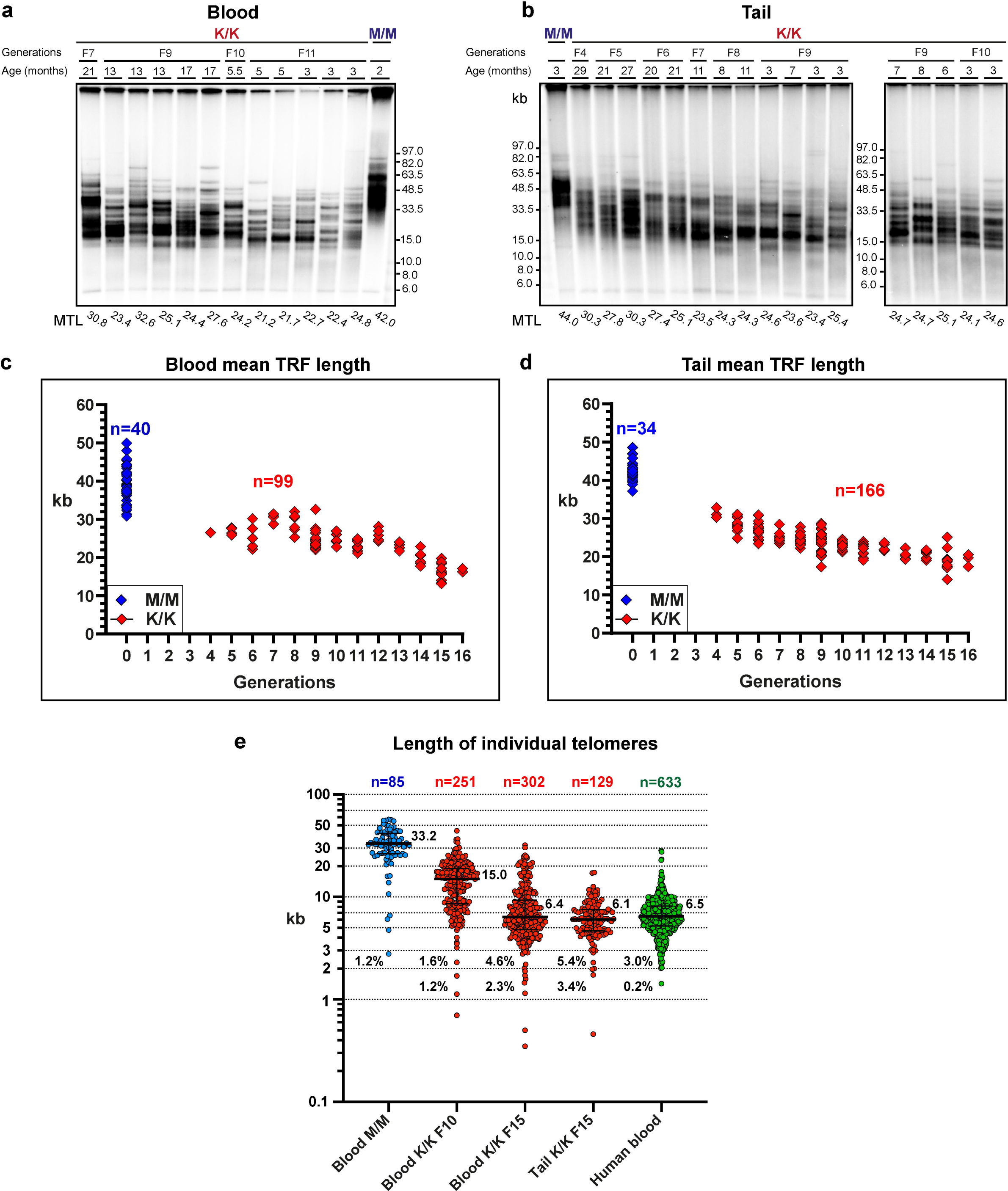
The Telomice telomeres progressively shorten over generations. Genomic DNA samples extracted from blood leukocytes (a) or tail (b) from mutant (K/K) or WT (M/M) mice at the indicated generations and ages were analyzed by PFGE and in-gel hybridization to the denatured DNA. Representative gels are shown. Mean TRF length for each generation was repeatedly measured from (c) blood (n= 40 for M/M and n= 99 for K/K) and (d) tail (n= 34 for M/M and n= 166 for K/K) samples in additional gels and plotted. The average age for K/K mice analyzed in the gels was 331 days, while the average age for M/M mice was 395 days. All data are summarized in Table S3. (e) Scatter plots show the length of individual telomeres in the indicated mouse (blood and tail) and human blood samples as determined by *NanoTelSeq*, with median and quartiles in horizontal lines. Median values in kb are indicated to the right of each scatter plot and the percentage of telomeres below 3 kb and 2 kb to the left of the plots. The details of all individual telomere reads are shown in Table S2.

*NanoTelSeq* also enabled us to determine the repeat lengths of specific mouse chromosome ends using the subtelomeric sequence to map them to the reference genome (Figure S11). We were only able to map reads to the ends of the long q-arm of the mouse acrocentric chromosomes since the subtelomere of the p-arm contains repetitive sequences that are not present in the reference genome. Our mapping revealed a large degree of heterogeneity in telomere lengths within and between chromosomes (Figure S11), the significance of which remains to be investigated in the future.

### Decreased regeneration potential of Telomice colonocytes *in vivo*

Telomerase null mice (*mTR^-/-^*) can be intercrossed for only six generations until fertility is lost^35^. Cells from the fourth *mTR^-/-^* generation (G_4_) onward frequently exhibit chromosome ends devoid of detectable telomere repeats, as well as end-to-end fusions and aneuploidy, correlating with defective spermatogenesis and reduced proliferation capacity of hematopoietic cells in the bone marrow and spleen^43^. In contrast, *Rtel1*^K/K^ Telomice breed readily until at least F16, indicating that telomeres in the germline maintain sufficient length and normal function. To examine if the short telomeres present in Telomice affect highly proliferative somatic tissues, we determined proliferation rates of intestinal stem and progenitor cells, which display the fastest proliferative rate of all cells in the body, by labeling of cells in S-phase using the thymidine analog EdU. We noted a small but significant decrease in the number of EdU-positive epithelial cells in the colon of Telomice (Figure 8a). To test if this proliferation defect is exacerbated under conditions of rapid epithelial renewal, we treated control and Telomice with dextran sodium sulfate (DSS), which causes ulcerative colitis-like pathologies due to its toxicity to colonic epithelial cells and engenders a strong regenerative response once the toxin is removed^44^ (see experimental outline in Figure 8b). Remarkably, while wild type mice showed the expected proliferative response, with ∼ 6 cells in S-phase per crypt cross section, the recovery was significantly blunted in Telomice, with only ∼ 3 replicating cells per crypt cross section (Figure 8c-e). These results suggested that while in the germline of Telomice telomerase is able to maintain sufficient (even if short) telomere length and normal cell proliferation, somatic cells are compromised in their ability to respond to situations requiring rapid regenerative proliferation, presumably because of their short telomeres and the absence of sufficient telomerase activity in somatic tissues.

**Figure 8.**
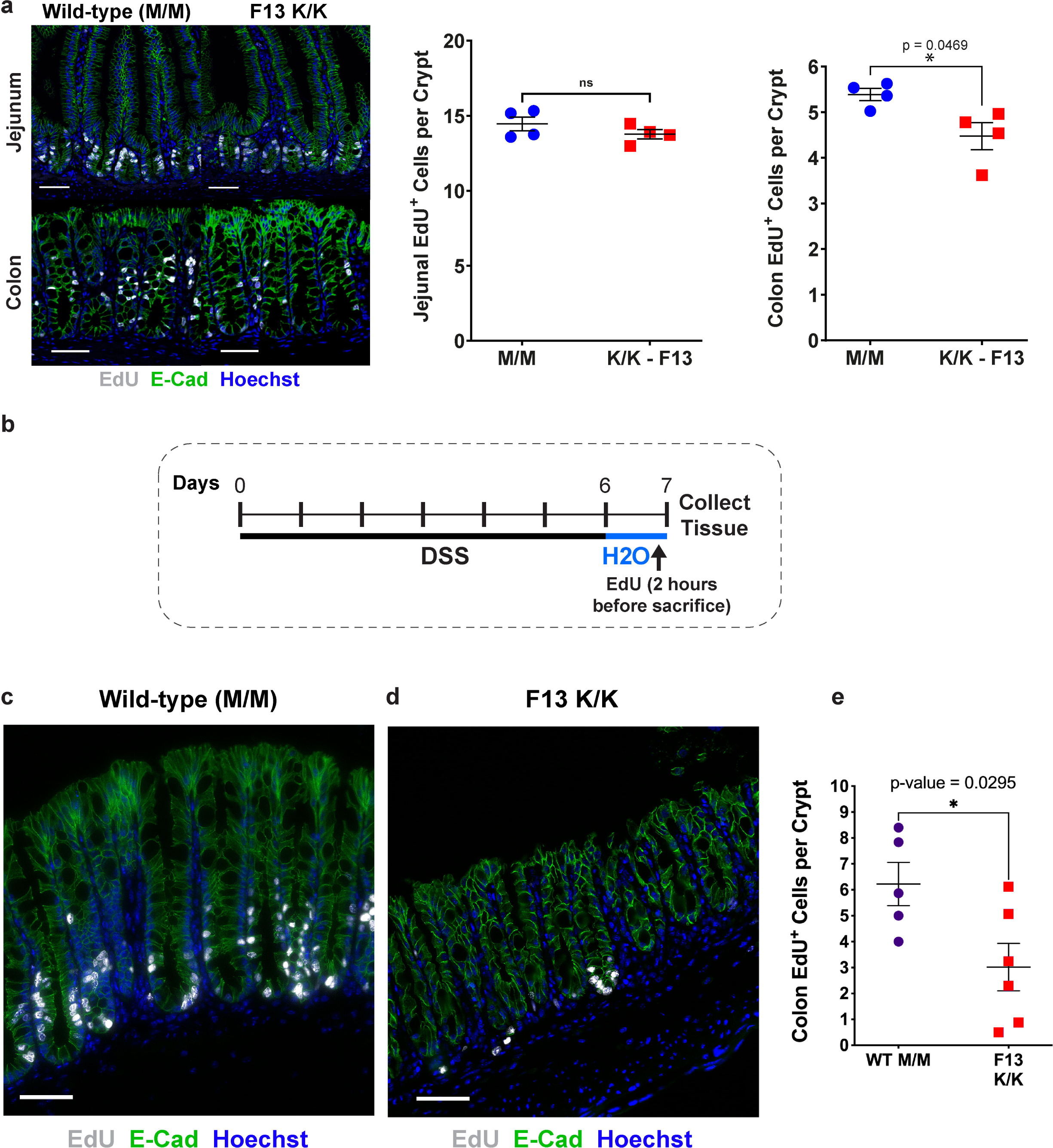
Intestinal regeneration following DSS-induced damage to colonocytes is impaired in Telomice. (a) Intestinal tissues from *Rtel1*^K/K^ mice (F13) were analyzed for the homeostatic proliferation rate in the gut epithelium by short-term (120 minutes) EdU labeling. Crypt stem and progenitor cells in the jejunum of mutant (K/K) mice displayed proliferation rates similar to those observed in control (M/M) mice. In the colon, however, K/K crypt cells divided somewhat less frequently. Purple circles, control (M/M); blue squares, mutant (K/K). Each data point represents one animal, with 10 crypts per animal counted and averaged into one data point (n = 4 for each group). Data are presented as mean +/-standard error of the mean, and significance was assessed using an unpaired, two-tailed Student’s t-test. Scale bars equal 50 μm. (b) Experimental timeline for dextran sulfate sodium (DSS) treatment. *Rtel1*^K/K^ mice (F13) and control (M/M) mice received 3% DSS in the drinking water for 144 hours, followed by 24 hours of regular drinking water to allow for colonic regeneration. Colons were collected and analyzed for crypt cell proliferation rate by short-term (120 minutes) EdU labeling. (c) Control (M/M) and (d) Rtel1K/K (F13) colons following DSS treatment were stained for EdU labeling (white), E-cadherin (E-Cad; green), and nuclei (Hoechst dye). Scale bars equal 50 μm. Note the reduced number of EdU-positive colonocytes in the Rtel1K/K mice (d). Scale bar equals 50 μm. (e) Quantification of replicating (EdU^+^) cells per crypt in DSS-treated WT (M/M) and *Rtel1*^K/K^ (F13) colons. Crypt cells in the colon of mutant (K/K) mice (n=6) displayed reduced proliferation compared to control (M/M) mice (n=5). Purple circles, control (M/M); blue squares, mutant (K/K). Each data point represents one animal, with 10 crypts per animal counted and averaged into one data point. Data are presented as mean and standard error of the mean, and significance was assessed using an unpaired, two-tailed Student’s t-test.

## DISCUSSION

RTEL1 was reported already in 2004 as a regulator of telomere elongation, based on its genetic association with telomere length in crosses between the long-telomere species *M. musculus* and the closely related, short-telomere species *M. spretus*^9^. However, the specific difference between the RTEL1 proteins from the two mouse species that is responsible for the different telomere length set points has remained unknown to date^12^. We derived a novel strain of *M. musculus* with a point mutation in *Rtel1*, replacing methionine 492 with a lysine, which established a new, *human* like telomere length set point, suggesting that this residue of RTEL1 is critical to its function in telomere length control. We termed this new *M. musculus* mouse strain ‘Telomouse’. All other mouse species examined had a methionine at position 492. Therefore the amino acid at this position cannot explain the short telomeres of other wild mouse species, which may rather result from another yet unknown variation in Rtel1 or in another telomere length regulator.

Following the growth of MEFs derived from an *Rtel1*^K/K^ embryo mouse over 250 population doublings and analyzing their telomeres by PFGE revealed that they gradually shortened to an averaged mean TRF length of 14.7 kb at PD 250, as compared to 37.8 kb of the WT MEFs at the same PD (Figure 2 and Table S1). However, detailed analysis of the TRF observed by PFGE and of qFISH hybridization signals, which correlates with the length of telomere repeat tracts, indicated that a large subtelomeric region (10-15 kb on average) is included in the TRF and the telomeric repeat tracts are highly heterogeneous in length (Figure 3), as proposed previously^35, 37^. Precise measurement of individual telomeres by Nanopore sequencing indeed revealed that the actual length of the telomeric repeat tracts is significantly shorter than the mean TRF length measured by PFGE (Figure 4). The *Rtel1*^K/K^ MEFs PD 250 telomeres had a median length of 5.6 kb as compared to 27.3 kb of WT MEFs at PD 250, consistent with a sub-telomere fraction of about 10 kb that is included in the TRF. Nevertheless, even the short telomeres of the *Rtel1*^K/K^ MEFs PD 250 largely retained their end-protection function, as indicated by their levels of genome-wide DDR foci, TIF, FISH aberrations and T-SCE, which were comparable to those in WT MEFs (Figure 5,6). Only telomere loss, indicated by a telomeric fluorescence hybridization signal below the detection limit, and Robertsonian fusions were elevated in the *Rtel1*^K/K^ MEFs PD 250 (Figure 6). Since these Robertsonian fusions did not exhibit a telomeric signal at the point of fusion, we assume that this tolerable type of fusion resulted from occasional shortening of telomeres below the critical length^45^. Interestingly, heterozygous *Rtel1*^M/K^ MEFs displayed telomeres nearly as short as the homozygous *Rtel1*^K/K^ MEFs (Figure 2a,b), suggesting that the *Rtel1*^M492K^ mutation represents a dominant separation-of-function mutation, which predominantly affects telomere length regulation but not other functions of RTEL1, consistent with the original discovery of RTEL1 as a dominant regulator of murine telomere length^9^.

Interbreeding Telomice over thirteen generations to F16 revealed that in each generation telomeres shortened until a new telomere length set point was reached. The observed rate of shortening, about 1 kb per generation, was much slower than the rate of shortening in the telomerase null (*mTR^-/-^*) mouse (4.8 kb per generation)^35^, indicating that telomerase action was not abolished but only reduced in Telomice. Importantly, telomeres did not shorten further from F15 to F16, in both blood or tail (Figures 7 and S8), suggesting that telomere length had stabilized, or was very close to stabilization, at a new telomere length set point in F15 and F16. Together, our results show that the M492K mutation altered the function of RTEL1 in telomere length regulation, in line with other reports suggesting that RTEL1 regulates telomerase action at telomere ends^9, 17, 26, 27, 30^. While we do not understand why the telomeres of *M. musculus* are ultra-long in the first place, the M492K mutation caused a dramatic reduction in the telomere length to a set point similar to that reached in *M. spretus* during natural evolution, without an apparent effect on the organismal health. This is in contrast to the disease-causing mutation in the same position of human RTEL1 (M492I), which caused genome-wide DNA damage and a fatal disease^17, 30, 42^.

To enable precise, base-resolution measurements of telomere length, we developed a novel method for vertebrate telomere length analysis by long read Nanopore sequencing. Nanopore sequencing has been used successfully for analyzing yeast telomeres^46^, but due to the different and variable yeast telomeric repeat, this method could not be employed for studying vertebrate telomeres and a different strategy needed to be developed for ligating the sequencing adapter to the telomere end of vertebrates. The long-read sequencing approach developed here provides the distribution, median telomere length, and, most importantly, the accurate length and abundance of the shortest telomeres, the latter of which cannot be determined by Southern blot, in-gel hybridization analysis, or qFISH. This is particularly important because the shortest telomeres are those dictating cell function and fate^47^. In addition, Nanopore sequencing can identify and assign telomere sequences to individual chromosome ends (Figure S11). Our Nanopore sequencing technique, termed *NanoTelSeq*, not only confirmed that the Telomouse telomeres are much shorter than those of WT *M. musculus* mice, but also enabled the precise analysis of the shortest telomeres not measurable by other methods. *NanoTelSeq* revealed a significant percentage of telomeres below 2 kb (but not below 1 kb) in Telomice MEFs, as well as in Telomouse blood and tail (Figures 4d and 7g). These observations suggest that the critical telomere length in both human and mouse is about 1 kb. Below this length telomeres cease to suppress DDR, arrest cell proliferation, and are depleted from the population of proliferating cells.

Short telomeres are major drivers of the aging-dependent decline in organ function and increase in cancer risk^5, 48^. Unfortunately, to date, these processes have been difficult to study in the common laboratory mouse, *M. musculus*, due to its extremely long telomeres. Attempts to study the implications of short telomeres in mice have thus far been based on telomerase null alleles, which cause ever-shortening telomeres and eventually result in severe pathologies and infertility^35^, manipulating telomere proteins to induce overt telomere deprotection^7, 49^, or the telomerase haploinsufficient *mTR^+/-^* strain of *M. musculus* crossed with the *M. castaneus* strain^50^. The resulting hybrid strain is not isogenic and is difficult to maintain because of the required heterozygocity with respect to both mTR and unknown telomere length regulators in the M. castaneus background. The Telomouse model derived here is genetically identical to the common *M. musculus* laboratory strain C57BL/6 except for the single amino acid difference in RTEL1. Yet, it displays dramatically shorter telomeres with no signs of telomere dysfunction under normal physiological conditions, with the exception of a small decrease in replication rate in the fastest proliferating cells of the colonic epithelium. However, when Telomouse colonocytes were challenged to mount a regenerative response, Telomouse epithelial cells were compromised in their replicative ability.

The utility of Telomouse depends in part on the ability to introduce and stably maintain additional genetic manipulations. While breeding of Telomice to another strain of mice would introduce a set of chromosomes with long telomeres and may require additional inter-crossing to reach short telomeres again, extensive backcrossing may not be needed if the short telomeres inherited from the Telomouse parent are maintained. The telomeres of the heterozygous M/K MEFs shortened almost in the same rate and extent as those of the homozygous K/K MEFs (Figure 2b) indicating that the M492K mutation is dominant, consistent with the original discovery of *Rtel1* as a dominant regulator of telomere length^9^. While this is yet to be examined in the mice, it is plausible that the short telomeres derived from *Rtel1*^K^ gametes in Telomouse crosses will not elongate in the heterozygous mice. In this case, the effects of manipulating an additional ‘gene X’ in the Telomouse would occur on the background of the short chromosomes inherited from the Telomouse parent. Alternatively, the genome of the Telomouse can of course be manipulated by CRISPR/Cas9 assisted genome editing in zygotes to introduce and examine the functional impact of additional loci, instead of crossing it to other strains with long telomeres. Therefore, we believe that the Telomouse is easier to maintain, manipulate and study than the unstable telomerase null or heterozygous strains. Altogether, the Telomouse model will open the way for a broad range of physiological experiments on the organismal level that were not possible previously, and reveal the effects of short telomeres in various tissues on cell proliferation, age-related organ decay, and cancer development.

## METHODS

### Generating a three-dimensional structure model of the murine RTEL1 helicase domain

We generated the murine RTEL1 (mRTEL1) 3D structure model consisting of residues 3-770 using the software Protein Homology/analogY Recognition Engine V 2.0 (Phyre 2), which produces a model of the protein of interest based on sequence alignment to known structures^31^. The 3D mRTEL1 structure was modeled using the cryo-EM structure of the yeast TFIIH helicase in the contracted state within the pre-initiation complex (PDB ID: 7O4K)^52^. TFIIH has 28% sequence identity to the mRTEL1 helicase domain. The presence of a structure with significant sequence identity with the murine gene enabled PHYRE to generate this model with 100% confidence. The model was further refined by applying geometry minimization in UCSF CHIMERA^53^. We then generated a model of RTEL1 in a complex with single stranded DNA, using the structure of the highly similar helicase domain of DinG (PDB ID: 6FWS), which belongs to the XPD family of helicases together with RTEL1, FANCJ and DDX11. The model was further refined by applying geometry minimization in Phenix^54^. The figures were generated in Pymol^55^.

### Derivation of Rtel1^M492K^ mice

CRISPR guide RNAs were designed as described^56^. For sgRNA preparation, the T7 promoter sequence was added to the sgRNA template by PCR amplification using pX335 (Addgene #42335) with Phusion high-fidelity DNA Polymerase Kit (NEB), Common sgRNA-R and Primer 1 or Primer 2 (see below).

Common sgRNA-R: 5’-AGCACCGACTCGGTGCCACT-3’

Primer 1 with gRNA sequence underlined:

5’-TTAATACGACTCACTATAGG**CATCTGCATCTCCAGAGCAA**gttttagagctagaaatagc-3’

Primer 2 with gRNA sequence underlined:

5’-TTAATACGACTCACTATAGG**CACCTGGAGGTCACAACACT**gttttagagctagaaatagc-3’

PCR products were purified using the QIAquick Gel Extraction kit (Qiagen), followed by *in vitro* transcription with the T7 High Yield RNA Synthesis kit (NEB). Newly synthesized sgRNAs were purified using the MEGAclear kit (Life Technologies), precipitated and diluted in injection buffer (10 mM Tris / 0.1 mM EDTA, pH 7.5) at a concentration of 500 ng/μl. The quality of the sgRNAs was confirmed using the total RNA Nano Chip on an Agilent bioanalyzer. The final injection mix was prepared using ssDNA repair template (IDT; 100 ng/μl), sgRNA1 (50 ng/μl), sgRNA2 (50 ng/μl), and Cas9 Nickase mRNA (Trilink; 100 ng/μl) in injection buffer. The sequence of the ssDNA repair template is shown below. The mutated A is underlined within codon 492 (in bold).

5’-GTTCGTACCCTTATCCTCACCAGCGGTACCCTGGCTCCACTGTCTTCCTTTGCTCTGGAG**AAG**CA GATGTATGTATGAGTCACCTGGAGGTCACAACACTAGGAACATGGTGGGTGGGGTTGG-3’

The final mix was spun twice at top speed for 30 minutes at 4°C to remove debris to avoid needle clogging. Cytoplasmic injection was done using C57Bl6 zygotes. After injection, SCR7 (Xcessbio) was added to the overnight egg culture at the concentration of 50 µM. Out of the 17 mice born, two had the targeted allele.

### Preparation and culture of MEF

Embryonic day (E)13.5 mouse embryos were dissected with head and inner organs removed, rinsed in HBSS, then digested with 300 μl papain isolation enzyme (ThermoFisher) for 30 minutes at 37°C following the manufacturer’s protocol. Embryonic digests were transferred to conical tubes with 1 ml of HBSS, pipetted up and down to achieve single cell suspension and then spun. The pellets were resuspended in MEF media (DMEM with 20% FCS, Pen/Strep/non essential amino acids) and plated. MEFs were grown in DMEM media containing 2 mM L-glutamine, penicillin-streptomycin, non-essential amino acids and 20% fetal bovine serum until immortalization (around PD60-70) and then with 10% fetal bovine serum in the same medium. Media and media supplements were purchased from Biological Industries Inc., Beit Haemek, Israel.

### Genomic DNA extraction

Leukocytes were obtained from blood by lysing red blood cells in 155 mM NH_4_Cl, 10 mM KHCO_3_ and 0.1 mM EDTA pH 8 followed by centrifugation. Leukocytes and MEFs were lysed in 10 mM Tris pH 7.5, 10 mM EDTA, 0.15 M NaCl, 0.5% SDS and 100 µg/ml proteinase K overnight at 37°C. Mouse tail samples were lysed in 100 mM Tris HCl pH 8.5, 5 mM EDTA, 0.1% SDS, 200 mM NaCl and 100 µg/ml proteinase K overnight at 50°C. Following cell lysis, high molecular weight genomic DNA was phenol-extracted, ethanol-precipitated, and dissolved in TE. Genomic DNA samples were examined by agarose gel electrophoresis of undigested DNA, and samples suspected to be degraded were excluded from further analysis.

### In-gel analysis of telomeric restriction fragments

Genomic DNA was treated with RNase A, digested with the *HinfI* restriction endonuclease, and quantified by Qubit fluorometer. Equal amounts (3-5 µg) of the digested DNA samples were separated by pulsed-field gel electrophoresis (PFGE) using a Bio-Rad CHEF DR-II apparatus in 1% agarose and 0.5 x TBE, at 14°C, 200V and pulse frequency gradient of 1 sec (initial) to 6 sec (final) for 20-22 hr (except for the K/K gel on the right in Figure 2a, which was electrophoresed in the same parameters but for 18 hr), or in regular gel apparatus, 0.7% agarose and 1 x TBE for 1,300 V x hr. Gels were ethidium-stained, dried and hybridized as previously described^30^. Probe mixtures contained a telomeric oligonucleotide (AACCCT)_3_, and mid range PFG ladder (NEB Inc.) and 1 kb ladder (GeneDirex Inc.), which were digested with *HinfI*, dephosphorylated by quick calf intestinal phosphatase (NEB Inc.) and heat-inactivated. All probes were 5’ end-labeled with [alpha-^32^P]-ATP and T4 polynucleotide kinase (NEB Inc.). The gels were exposed to a PhosphorImager. TRF length was quantified using *TeloTool* (corrected mode)^51^. Individual band lengths were interpolated from the regression formulas for plotting the log of the ladder band sizes as a function of the migration distance, and native and denatured in-gel hybridization intensity was quantified using *ImageQuant-TL* (GE Healthcare Inc.).

### Immunofluorescence of cultured cells

Cells were seeded onto glass coverslips and grown for 1-4 days. Immunofluorescence was performed as described^51^ with the following changes: Primary antibodies used were TRF1 (Abcam; AB10579) and γH2AX (Cell Signaling Technology; 2577S), incubated overnight at 4°C. Imaging acquisition was by FV-1200 confocal microscope (Olympus, Japan) with a 60X/1.42 oil immersion objective. *NIH ImageJ*^38^ was used for image analysis and foci quantification.

### Metaphase FISH

Metaphase spread and FISH were performed as described^30^, with the following changes: Cells were arrested in metaphase by 3-4 hr incubation with colcemid. After fixation, both MEFs and human fibroblasts were mixed and dropped on the slides. The PNA probes used were TelC-Alexa488 telomeric probe (F1004, Panagene Inc.) and CENPB-Cy3 centromeric probe (F3002, Panagene, Inc). Imaging acquisition was by FV-1200 confocal microscope (Olympus, Japan) or by a Nikon Eclipse Ti-E microscope equipped with a CoolSNAP Myo CCD camera. The telomeric signal was quantified using *Telometer*^57^.

### Meta-TIF assay

We followed the protocol developed by Cesare and colleagues^58^ with a few changes. MEFs were synchronized in metaphase with 0.2 μg/ml colcemid for 1-2 hrs, collected by trypsinization, and re-suspended in fresh hypotonic solution (0.2% KCl and 0.2% Tri-sodium citrate in DDW) at room temperature (RT). 500 μl of swollen MEFs (200,000 cells/ml) were pipetted into each cytocentrifuge (Thermo Scientific Cytospin 4) funnel and centrifuged for 7-10 min at 1,000 rpm with medium acceleration rate onto Superfrost glass slides. The slides were immediately fixed in 1x PBS / 4% formaldehyde at RT for 10 min, permeabilized in KCM buffer (120 mM KCl, 20 mM NaCl, 10 mM Tris pH 7.5 and 0.1% Triton X-100) at RT for 10 min, and then blocked with antibody dilution buffer (20 mM Tris pH 7.5, 2% BSA, 0.2% fish gelatin, 150 mM NaCl, 0.1% Triton X-100 and 0.1% sodium azide) containing DNase-free RNase A for 15 min at 37°C. The slides were incubated overnight at 4°C, with the primary antibody (γH2AX [Millipore; JBW 301]) diluted 1:1000 in antibody dilution buffer. After washing, the slides were incubated with a secondary antibody (Thermo Scientific, A-11029) diluted 1:500 in antibody dilution buffer for 30 min at RT, washed and fixed for 10 min in 1x PBS with 4% formaldehyde at RT. The slides were dehydrated using a graded ethanol series (70% for 3 min, 90% for 2 min and 100% for 2 min) and then air-dried. FISH was performed using the metaphase FISH protocol described above, but with denaturation at 80°C for 5 minutes, hybridization overnight at RT using the TelC-Cy3 probe (F1002-5, Panagene), and washing in PNA wash A (70% formamide and 10 mM Tris pH 7.5), then in PNA wash B (50 mM Tris pH 7.5, 150 mM NaCl and 0.08% Tween-20). After rinsing several times with dH_2_O, the slides were mounted with Vectashield + DAPI (ThermoFisher Scientific Inc.) and stored at least overnight before imaging.

### Chromosome-Orientation Fluorescent In-Situ Hybridization (CO-FISH)

MEFs were cultured in medium supplemented with 7.5 μM bromodeoxyuridine (BrdU) and 2.5 μM bromodeoxycytidine (BrdC) for 10 hours, and metaphase spreads were prepared as in the metaphase FISH assay, adding Colcemid (without changing the medium), and incubating for 3 hours. Slides were treated with RNase A for 30 minutes at 37°C, stained with Hoechst 33258 (0.5 μg/ml in 2x SSC) for 30 minutes at RT, irradiated with 365-nm UV light for 5.4×103 J/m2, and then digested with Exonuclease III (9.5 μl of 100 units/µl) at 37°C for 30 minutes. Slides were washed in 1x PBS, dehydrated in ice-cold ethanol series of 70%, 85% and 100% for 2 min each and air-dried in the dark. For PNA hybridization, the same hybridization mix as in the metaphase FISH assay was used, hybridizing 1 hour with the TelG probe at RT in the dark, rinsing in wash I (10 mM Tris-HCl pH 7.2, 70% Formamide and 0.1% BSA in 1x PBS) for 2 minutes, drying briefly by dabbing, then hybridizing for another 1 hour with the TelC probe at RT in the dark. After removing the coverslips in 1x PBS, the slides were washed twice in wash I for 15 min each time, then in wash II (0.1 M Tris-HCl pH 7.2, 0.15 M NaCl, 0.08% Tween-20) three times for 5 min each time. Finally, the slides were washed in 1x PBS three times, 5 min each, dehydrated in ethanol baths (70%, 95%), air dried in the dark, and mounted as in the meta TIF assay.

### Nanopore sequencing of telomeres

Single samples were sequenced separately using six telorette oligonucleotides, each ending with a different permutation of the six base telomeric repeat at its 3’ end (Figure 4a). 10-20 µg of high molecular weight genomic DNA, prepared as described above and treated with RNase A, were ligated overnight at 20°C with a mix of six 5’ phosphorylated telorette oligonucleotides (0.1 µM each) and 2,000 units of T4 ligase (NEB Inc.) in a 50 µl reaction. The ligated DNA was then annealed to 1µM teltail-tether oligonucleotide by heating for 10 minutes at 65°C and cooling slowly to room temperature. The DNA was purified using AMPure XP beads (Beckman Coulter Life Sciences), and ligated to the Nanopore sequencing adapter (AMII) following the protocol for genomic DNA ligation sequencing with native barcoding (SQK-LSK109 with EXP-NBD104; Oxford Nanopore Technologies). For multiplexed reactions, 2-6 samples were pooled in a sequencing library using barcoded telorette 3 oligonucleotides. The barcoded telorette oligonucleotides were first annealed to a complementary teltail-tether oligonucleotide and then each double-stranded telorette with a specific barcode (1 µM) was ligated to an individual DNA sample (1-10 µg) as before. The ligation reactions were stopped by adding 20 µM EDTA and the barcoded samples were combined, purified, ligated to 3 µl sequencing adapter (AMII) and sequenced following following the standard protocol for genomic DNA ligation sequencing with native barcoding (SQK-LSK109 with EXP-NBD104; Oxford Nanopore Technologies), except for the library that included the K/K mouse blood sample, which was purified by polyethylene glycol (PEG) precipitation instead of AMPure XP beads following^59^. 0.4-5.5µg of the Sequencing libraries were loaded onto a Nanopore R9 flow cell. The sequencing library purified by PEG was loaded with SQB buffer (Oxford Nanopore Technologies) without loading beads due to the high viscosity of the library.

### Oligonucleotides used for sequencing. Underlined is the telomeric repeat. P, 5’ phosphate

Telorette1: 5’-P-TGCTCCGTGCATCTCCAAGGTTCCTAAC-3’ Telorette2: 5’-P-TGCTCCGTGCATCTCCAAGGTTAACCCT-3’ Telorette3: 5’-P-TGCTCCGTGCATCTCCAAGGTTCTAACC-3’ Telorette4: 5’-P-TGCTCCGTGCATCTCCAAGGTTTAACCC-3’ Telorette5: 5’-P-TGCTCCGTGCATCTCCAAGGTTACCCTA-3’ Telorette6: 5’-P-TGCTCCGTGCATCTCCAAGGTTCCCTAA-3’

Telorette3-NB01n:

P-5’-TGCTCCGTGCATCTCC-AAGGTTAA-CACAAAGACACCGACAACTTTCTT-CAGCACCT-CTAACC-3’

Telorette3-NB02n:

P-5’-TGCTCCGTGCATCTCC-AAGGTTAA-ACAGACGACTACAAACGGAATCGA-CAGCACCT-CTAACC-3’

Telorette3-NB03n:

P-5’-TGCTCCGTGCATCTCC-AAGGTTAA-CCTGGTAACTGGGACACAAGACTC-CAGCACCT-CTAACC-3’

Telorette3-NB04n:

P-5’-TGCTCCGTGCATCTCC-AAGGTTAA-TAGGGAAACACGATAGAATCCGAA-CAGCACCT-CTAACC-3’

Telorette3-NB07n:

P-5’-TGCTCCGTGCATCTCC-AAGGTTAA-AAGGATTCATTCCCACGGTAACAC-CAGCACCT-CTAACC-3’

Telorette3-NB08n:

P-5’-TGCTCCGTGCATCTCC-AAGGTTAA-ACGTAACTTGGTTTGTTCCCTGAA-CAGCACCT-CTAACC-3’

Teltail-tether:

5’-AACCTTGGAGATGCACGGAGCAAGCAAT-3’

### Computational processing of Nanopore sequencing reads

Deducing the nucleotide sequence (base calling) from the raw data (Fast5 files) was performed with the Nanopore *Guppy* application at high or super accuracy mode. The sequences (FastQ files) were filtered and analyzed by a dedicated application termed *NanoTel*, which is available on GitHub. Since we noticed a difficulty of the base calling application in distinguishing between purines (A and G) in the context of the telomeric repeats, the repeats on the C-strand were identified by searching for the sequence CCCTRR (R represents a purine) in the six possible permutations. The telomere length and read length were extracted and presented on plots of telomere density along the entire read. Telomere density was calculated in a moving window of 100 nt and describes the portion within each 100 nt sequence that contains telomeric repeats (from 0 - no telomeric sequence to 1 - fully telomeric). A telomere is identified by a summed telomere density of at least 2 for at least three consecutive windows, each having a telomere density of at least 0.3. Then, the telomere beginning and end are defined by the first and last telomeric repeat within the identified windows or flanking windows.

### Mapping of telomere reads to specific chromosome ends

The subtelomeric sequences of telomeric reads were mapped to the *M. musculus* GRCm39 reference genome using the *minimap2* program with the recommended set of parameters optimized for long Nanopore sequencing reads^60^. Alignment results were filtered according to the following criteria: (1) At minimum of 70-90% (depending on the sample) of the read’s subtelomeric region was aligned. (2) The alignment started within the most distal 10,000 nt of a chromosome end in the reference genome. (3) The read’s subtelomere was aligned to the reference sequence in the correct orientation with respect to the chromosome end.

### Selection of telomere reads for median calculation

Reads with an overall telomere density of 0.7 were analyzed. Since not all reads were sufficiently long to exclude a bias for short telomeres, reads were further selected based on their length. Telomere reads were sorted for sequence length, from long to short, and a running median for telomere length was calculated from the longest read down to each point. Only reads longer than the median telomere length were selected. These reads were used for calculating the median telomere length for each sample. For the reads mapped to chromosome ends, only reads longer than the median + 1,000 nt were chosen.

### DSS treatment

Dextran sulfate sodium (DSS) (MP Biomedicals, cat. # 0216011001) was administered to Rtel1K/K mice (F13) and WT (M/M) mice via drinking water (3% DSS) for 144 hours. All mice received regular drinking water for 24 hours prior to sacrifice.

### Tissue Immunohistochemistry and Immunofluorescence

For EdU treatment, 150 μl of 10 mg/ml EdU solution was injected intraperitoneally two hours before mice were sacrificed. After euthanasia, mouse tissues were rinsed in PBS and fixed with 4% paraformaldehyde overnight. Fixed tissue was rinsed in PBS three times for 30 minutes per rinse and dehydrated for paraffin embedding and sectioning. EdU was detected according to the manufacturer’s protocol (Click-iT Plus EdU Cell Proliferation Kit for Imaging, Alexa Fluor 647 dye, C10640, Invitrogen). Briefly, tissue sections were deparaffinized, and heat antigen retrieval was performed in a pressure cooker with Citrate Buffer, pH 6 (64142-07, Electron Microscopy Sciences). The tissue sections were then incubated in the Click-iT Edu Reaction Cocktail for 30 minutes at room temperature, washed, and blocked with CAS-Block (008120, Invitrogen) for 30 minutes at room temperature. Tissue sections were stained overnight with a mouse anti-E-Cadherin (610181, 1:500, BD Biosciences) primary antibody. The following day, tissue sections were rinsed in PBS and incubated for two hours at room temperature with a Cy2 anti-mouse secondary antibody. The tissue sections were washed in PBS and stained with Hoechst (H3570, 1:10,000, Molecular Probes) to visualize nuclei. Images were taken on a Leica laser scanning confocal microscope (TCS SP8, Penn Cell & Developmental Biology Microscopy Core). Images were processed and brightness and contrast were enhanced using FIJI (*ImageJ*, Version 2.1.0, NIH)^38^.

### Mice

DNA from a total of 120 K/K and M/M (100% pure Blk6) mice were analyzed multiple times in separate gels (as detailed in Table S3). The average age for K/K mice analyzed in the gels was 331 days, while the average age for M/M mice was 395 days. At each specific generation of breeding, the following mice were analyzed:

F0 (M/M): 24 mice (63 – 665) days old, F4 (K/K): one mouse 881 days old,

F5 (K/K): 7 mice (341 – 895) days old,

F6 (K/K): 13 mice (318 – 787) days old,

F7(K/K): 2 mice (329 – 872) days old, F8 (K/K): 7 mice (233-778) days old,

F9 (K/K): 28 mice (81-402) days old, F10 (K/K): 7 mice (42-68) days old, F11 (K/K): 9 mice (68 - 138) days old,

F12 (K/K): 9 mice 118 and 171 days old, F13 (K/K): 9 mice: 80 days old,

F14 (K/K): 4 mice: 186 and 205 days old,

F15 (K/K): 8 mice: 76, 95, and 182 days old,

F16 (K/K): 2 mice: 144 and 163 days old.

Three mouse embryonic fibroblast cultures were grown over 250 population doublings (PD) and sampled every 10 PD (a total of 121 samples). Samples were analyzed in multiple gels, as detailed in Supplementary Table S1.

## Data availability

Representative gels are shown in Figures 2, 7, S2-S4, and S7-S10. All measured and calculated values are provided in the Tables S1 (MEFs) and S3 (mice). The complete un-spliced gel images for Figure 2a are shown in Figure S2a,c,e. The complete un-spliced gel images for Figure 7b, including the native hybridization, are shown in Figure S9d,h. Additional gel replicates and data are available upon request to the corresponding authors.

The nanopore telomeric read characteristics are summarized in Table S2.

### Statistical analysis

Statistical analysis was performed by Microsoft Excel and GraphPad Prism 8.0, using simple linear or segmental linear regression and setting the break points manually to give the best fit by coefficient of determination (R^2^). For statistical tests two-tailed paired and unpaired t-test, nested t-test, and nested one way ANOVA were used, as indicated for each figure. P < 0.05 was considered statistically significant.

## ACKNOWLEDGEMENTS

This research was supported in parts by the Israel Science Foundation grant 2071/18 to Y.T., the NIDDK through R37-DK053839 and R01-CA249929 to K.H.K., and by the Israel-UK-Palestine GROWTH Fellowship to R.S. We are grateful to all members of the Tzfati and Kaestner laboratories for stimulating discussions and assistance. We thank Ittai Ben-Porath and Amir Eden for the gifts of WT mice and MEFs; Devora Olam for assistance in mouse blood collection; Naomi Melamed-Book, Bio-Imaging Unit at The Hebrew University and Manish Goyal for assistance with confocal imaging; Haimasree Bhattacharya, the Center for Interdisciplinary Data Science Research (CIDR), The Hebrew University; Dana Avrahami-Tzfati for assistance with CRISPR; and Ran Avrahami for assistance with statistical analyses. We thank the University of Pennsylvania’s Diabetes Research Center (DRC) for the use of the Functional Genomics Core (P30-DK019525) and the Center for Molecular Studies in Digestive and Liver Diseases for the use of the Molecular Pathology and Imaging and Transgenic and Chimeric Mouse Cores (P30-DK050306). We also thank Lan Cheng in IDOM Histology Service for performing the IF staining, and Ayano Kondo for writing a code for *ImageJ* analysis.

## AUTHOR CONTRIBUTIONS

Corresponding authors Y.T. (telomere biology) and K.H.K. (mouse biology): study concept and design; analysis and interpretation of data; writing and revising the manuscript. Y.T. R.S.: study concept and design; analysis of telomere phenotypes; interpretation of data; writing and revising the manuscript. C.L.M. and H.K.: derivation and *in vivo* analysis of the mouse strains, and writing and revising the manuscript. N.E.: analysis of telomere phenotypes and computational analysis of Nanopore sequencing data. D.L.: computational analysis of Nanopore sequencing data. E.S.: computational analysis of the mouse RTEL1 structure. Y.R., A.M., V.O. and M.T.: *in vivo* analysis of the mouse strains.

