## Supplementary Information for "Telomouse – a mouse model with human-length telomeres generated by a single amino acid change in RTEL1"

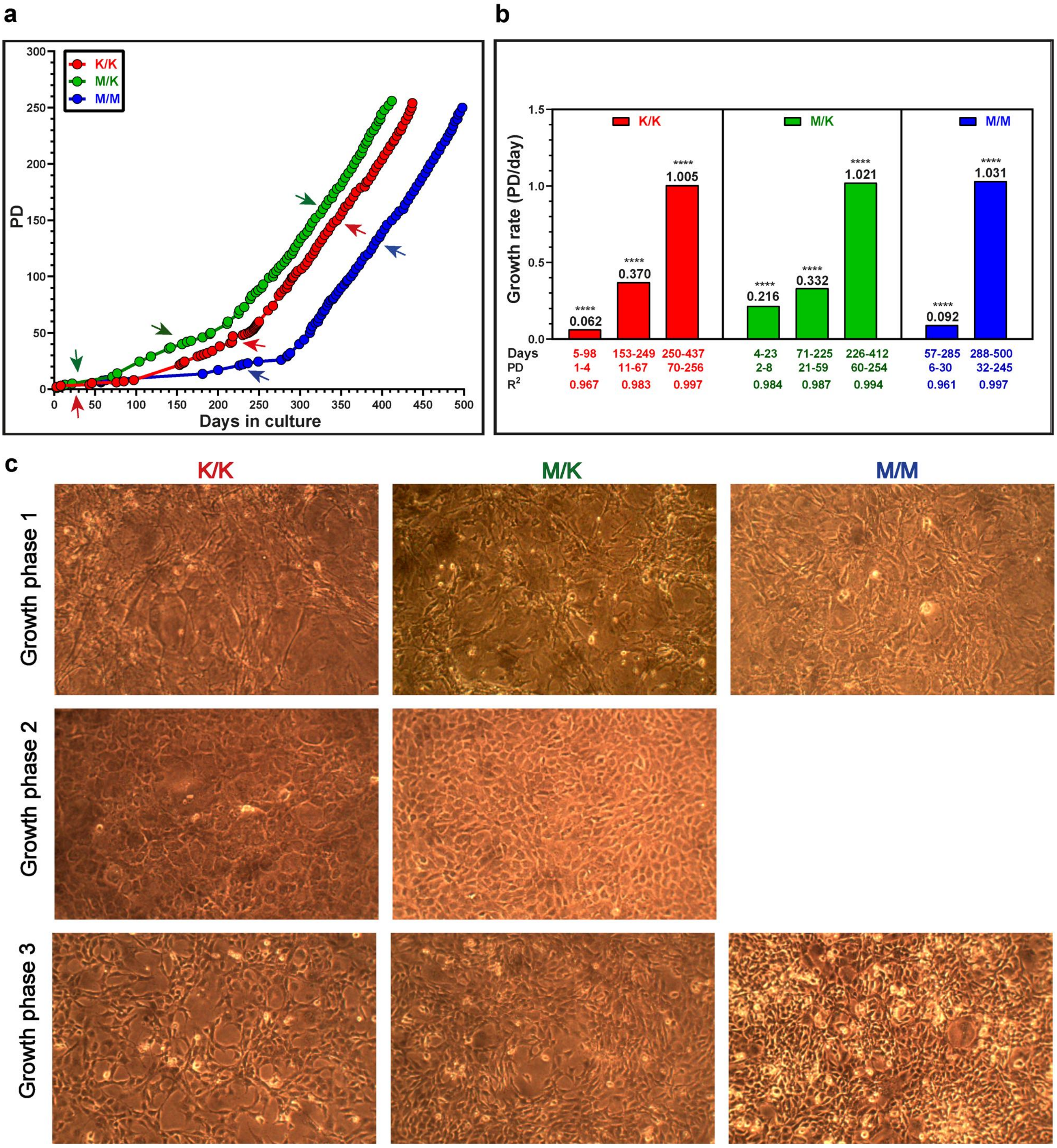


**Figure S1. Immortalization and growth characteristics of MEFs from WT and *Rtel1*$^{M492K}$ mutant mice.** (a) MEF cultures were prepared from F3 littermate embryos (K/K and M/K) and from WT *Mus musculus* embryos (M/M), immortalized by serial passaging and grown to PD 250. Arrows indicate different growth phases. (b) Growth rates were calculated based on the graphs in (a) and indicated above the bars. Days, PD, and coefficient of determination ($R^2$), for the linear regression line calculated for each growth phase are indicated below. **** P < 0.0001. (c) Representative images show different cell morphology for each growth phase.

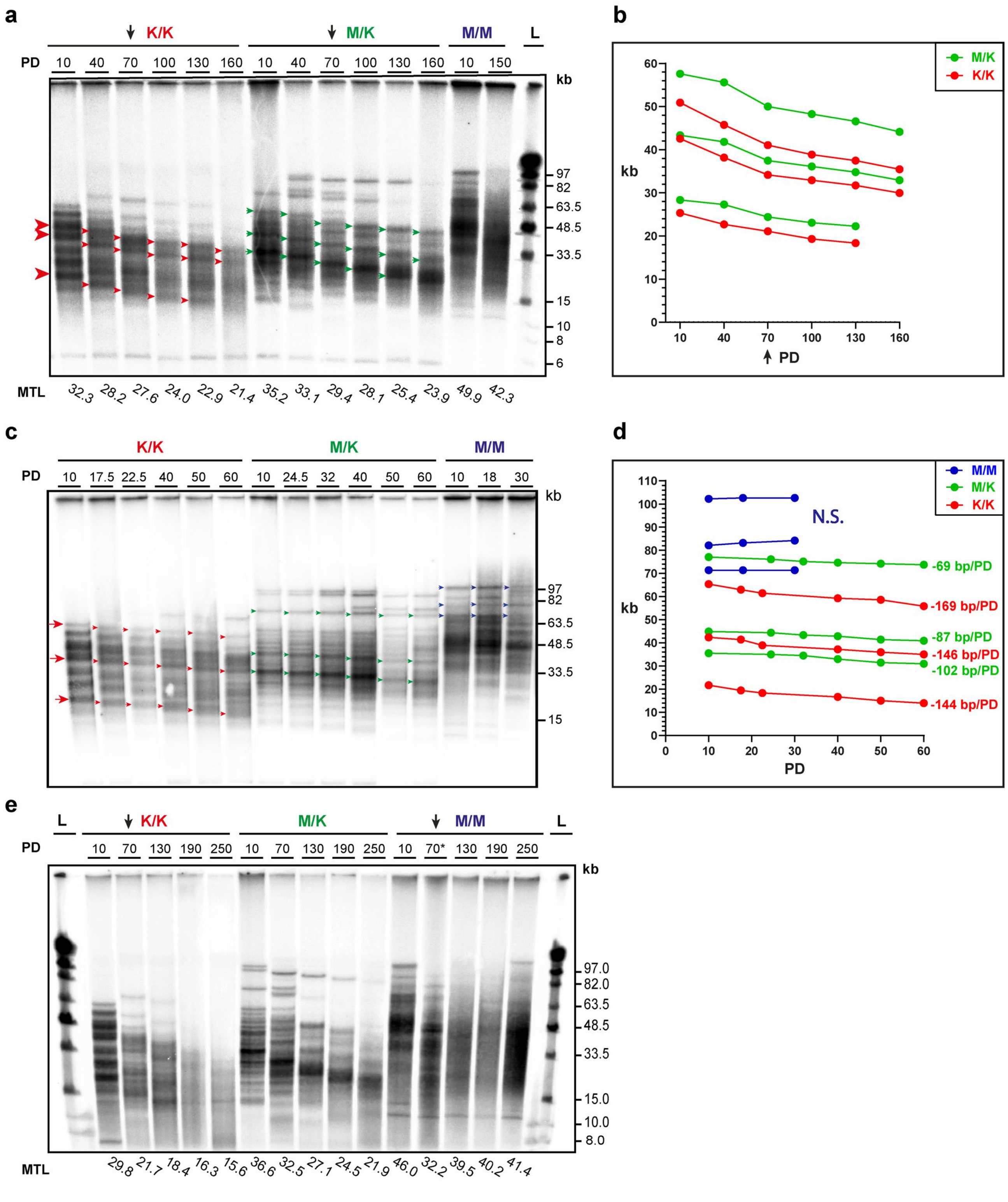


**Figure S2. MEF telomeres display a distinct banding pattern.** (a,c,e) Genomic DNA samples prepared from *Rtel1*$^{M492K}$ homozygous (K/K) or heterozygous (M/K) mutant or WT (M/M) MEF cultures were analyzed by PFGE and in-gel hybridization. The complete change of cell morphology (see Figure S1c), reflecting cellular immortalization, is indicated by arrows on the gel images. M/M** represents a MEF culture derived from a progeny of two heterozygous RTEL1 mutant mice. Since this embryo inherited short telomeres it is not considered as 'true' WT and thus was excluded from further analysis. M/M PD 160* and 250* DNA samples were suspected to be degraded based on PFGE of uncut DNA, and were also excluded. (b,d,f) The length of distinct TRF bands indicated by small arrows on the gel images was calculated and plotted. The rate of shortening is indicated. 'N.S.', no significant deviation from a horizontal line with a slope of 0.

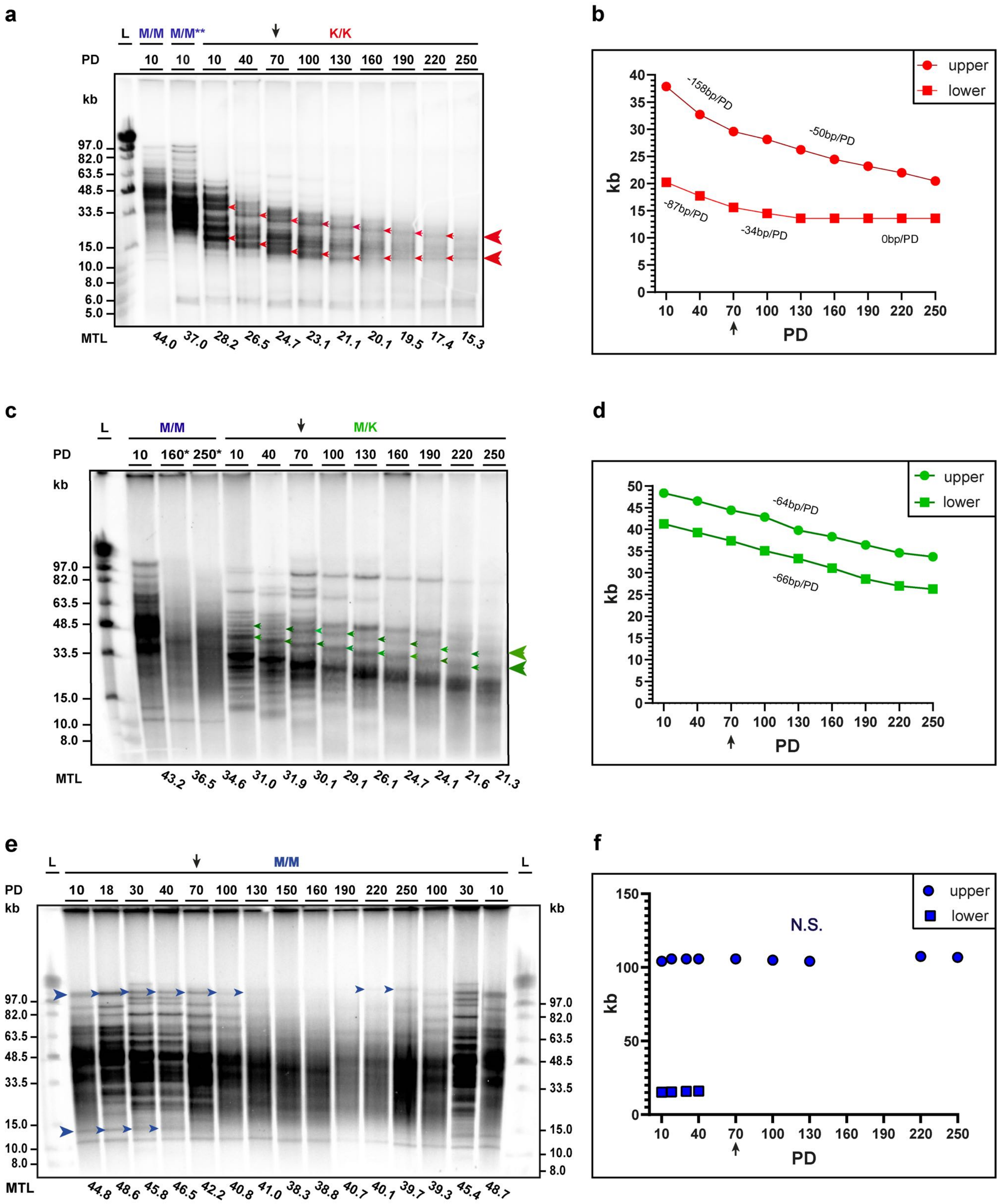


**Figure S3. MEF telomeres display a distinct banding pattern.** (a,c,e) Genomic DNA samples prepared from *Rtel1*$^{M492K}$ homozygous (K/K), heterozygous (M/K), or WT (M/M) MEF cultures were analyzed by PFGE and in-gel hybridization. The complete change of cell morphology (see Figure S1c), reflecting cellular immortalization, is indicated by arrows on the gel images (a,e). The M/M sample PD 70* in (e) was suspected to be degraded and excluded. (b,d) The length of distinct TRF bands indicated by small arrows on the gel images was calculated and plotted. The rate of shortening is indicated. 'N.S.', no significant deviation from a horizontal line with a slope of 0.

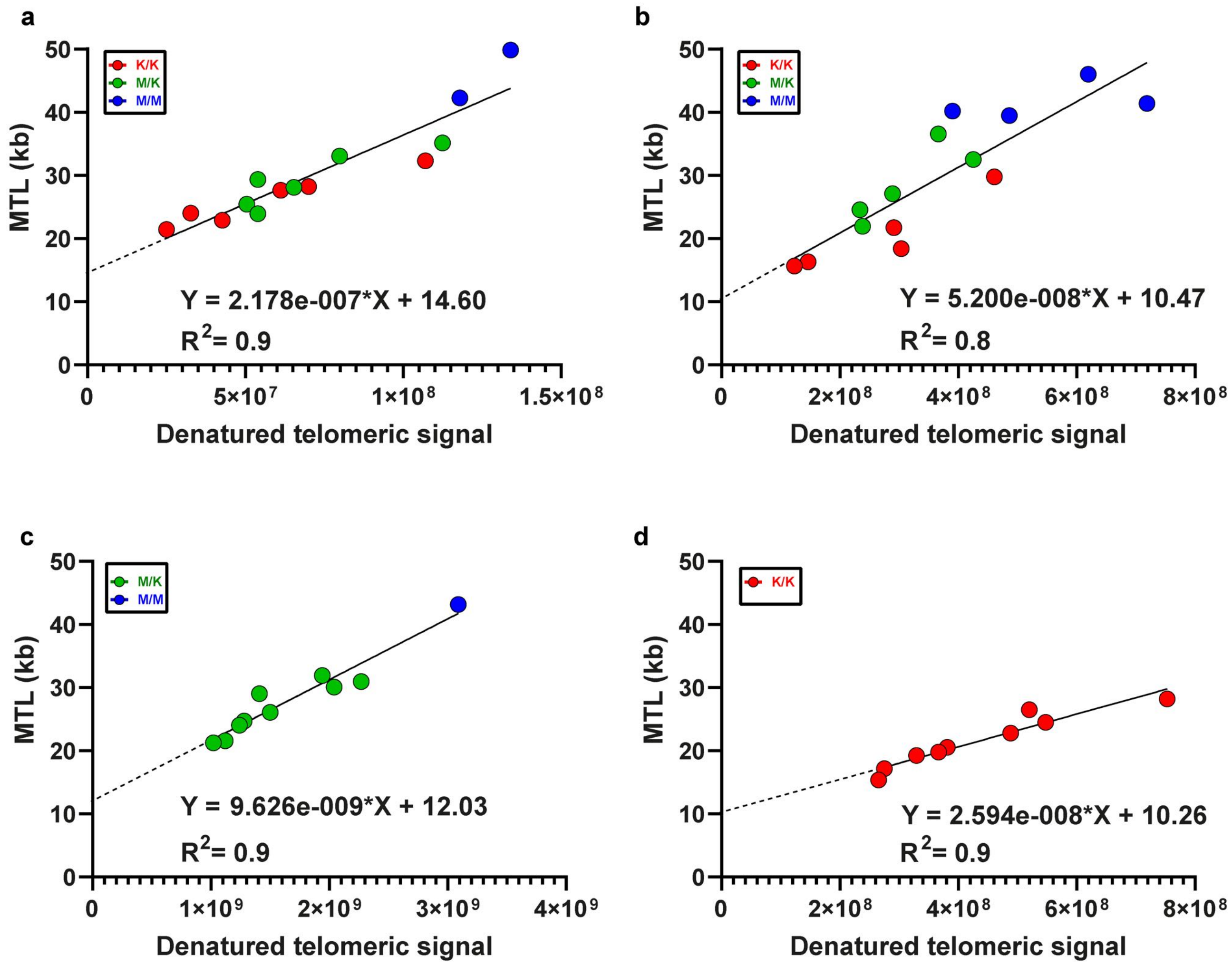


**Figure S4. Telomeric repeat arrays are shorter than the telomeric restriction fragments.** Mean TRF length was plotted as a function of the denatured hybridization signals quantified by *ImageQuant-TL* for the gel shown in Figure S3a (a), Figure S3e (b), Figure S2c (c), and Figure S2a (d). The linear regression line, formula and coefficient of determination ($R^2$) value are shown for each graph. Blue indicates M/M (WT), green M/K (heterozygous); and red indicates K/K (homozygous) samples.

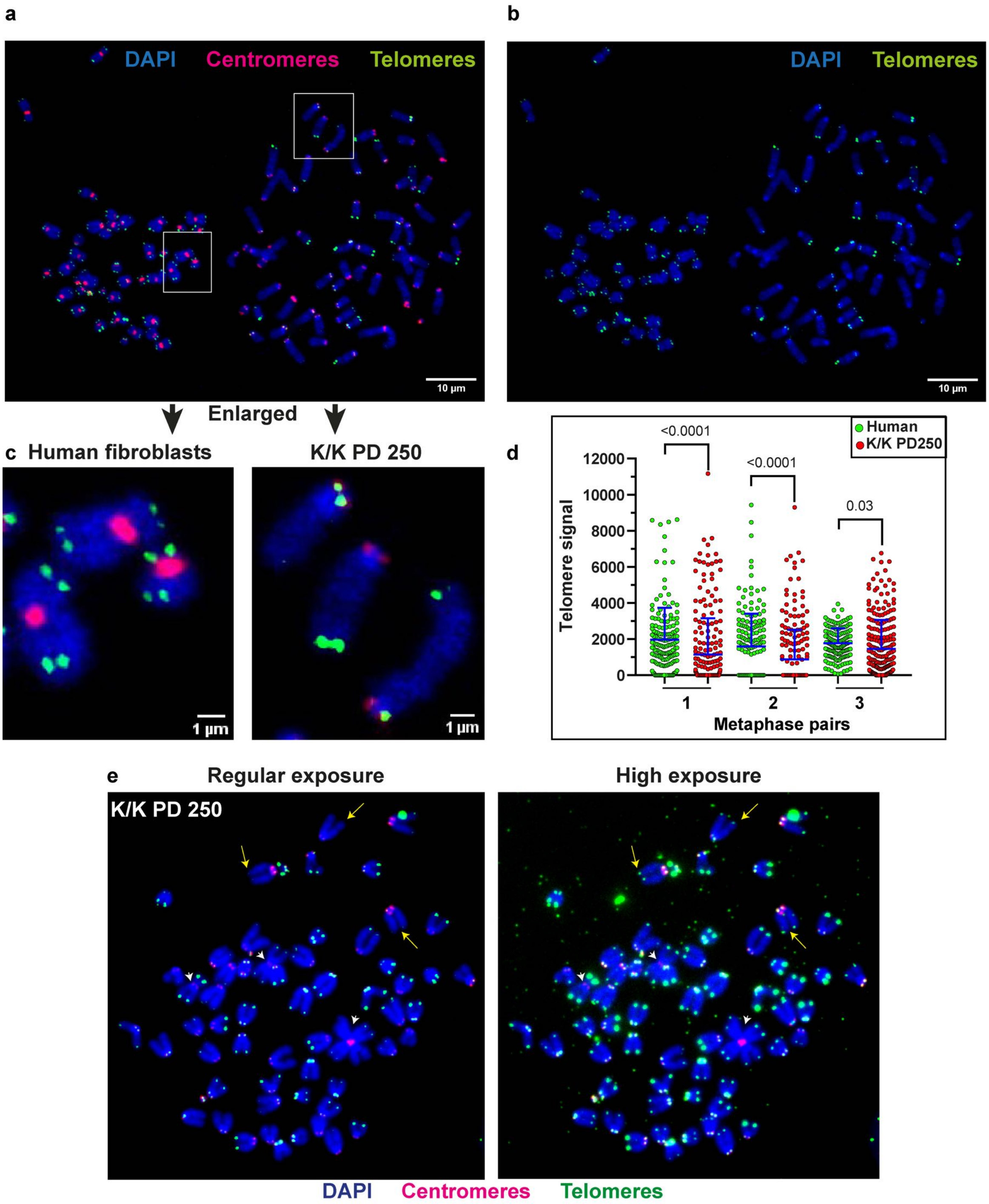


**Figure S5. *Rtel1*$^{K/K}$ MEFs telomeres are more heterogeneous than human fibroblasts telomeres.** (a) *Rtel1*$^{K/K}$ MEFs at PD 250 and human telomerase positive fibroblasts with a mean TRF length of 14 kb (as measured by in-gel hybridization; Figure 3d) were arrested in metaphase, mixed and spread on slides. An example of one microscope field is shown. The slides were hybridized to a green telomeric PNA probe and a red centromeric PNA probe to distinguish the mostly metacentric human chromosomes from the mouse acrocentric chromosomes. Imaging acquisition was by FV-1200 confocal microscope (Olympus, Japan). (b) Shows the same image without the centromeric signal and (c), enlargements, showing more undetected telomeric signals in K/K MEFs than human fibroblasts. (d) The telomeric signals were quantified for three pairs of human and mouse metaphases. 164 - 172 human and 236 - 244 mouse chromosome ends were quantified in each image using the *Telometer* plugin of *NIH ImageJ*[38] and plotted. Horizontal lines indicate the mean and standard deviation. The P values were calculated by a 2-tailed unpaired t-test. (e) High exposure (right) reveals very short telomeres (long arrows) in KK PD 250 MEFs undetected by standard exposure (left). Telomere loss under high exposure is associated with chromosome fusion (short arrows).

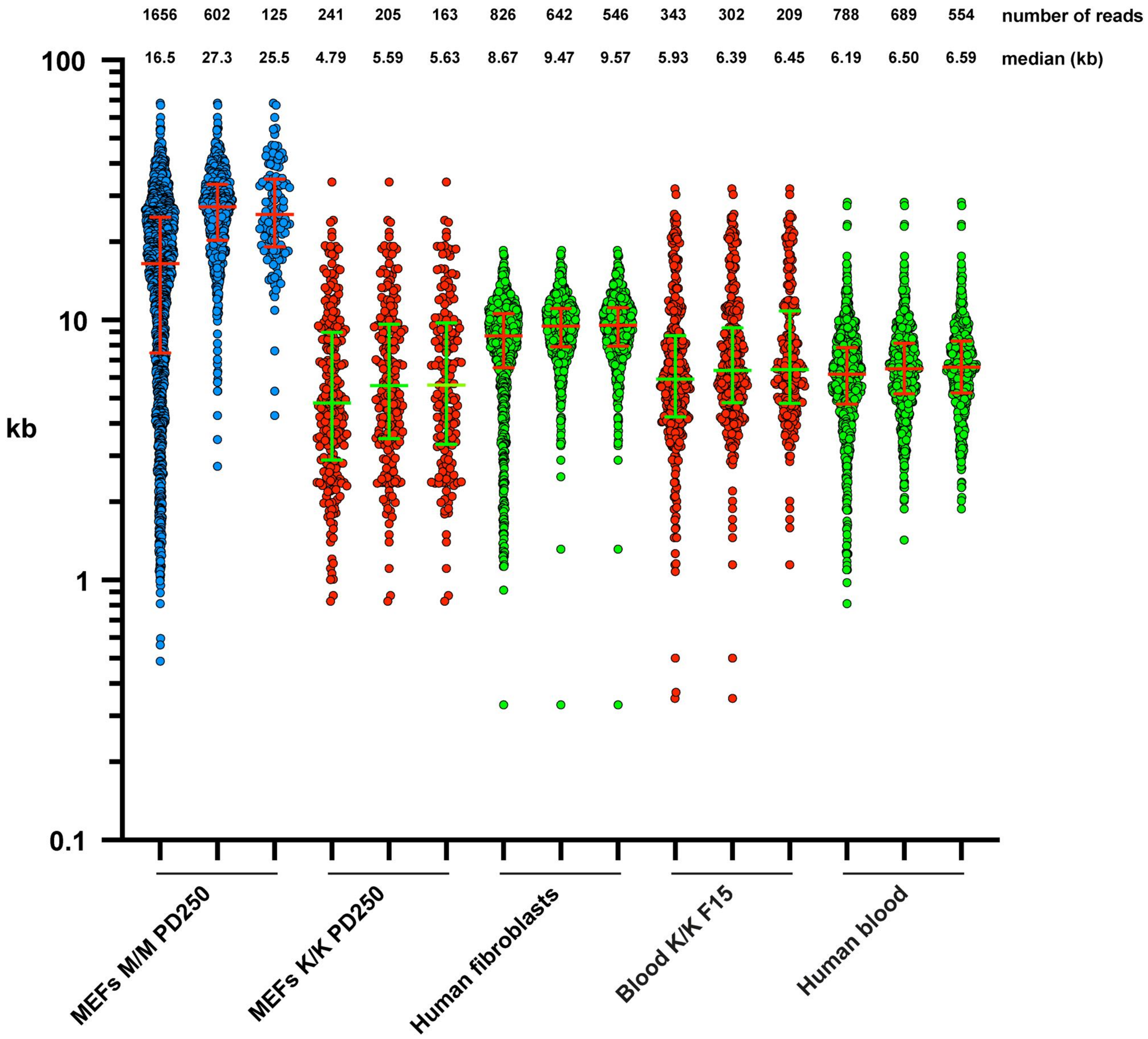


**Figure S6. Filtration of telomere reads obtained by Nanopore sequencing based on read length.** The length of single telomeres as calculated by *NanoTel* is shown on a scatter plot. For each sample, the plot on the left shows all telomeric reads, the middle plot shows telomeres with an overall read length longer than the median telomere length, and the right plot shows telomeres longer than the longest incomplete telomere (see Materials and Methods). Horizontal lines indicate the median and quartiles. The number of telomeric reads and the calculated median values are shown above the plots.

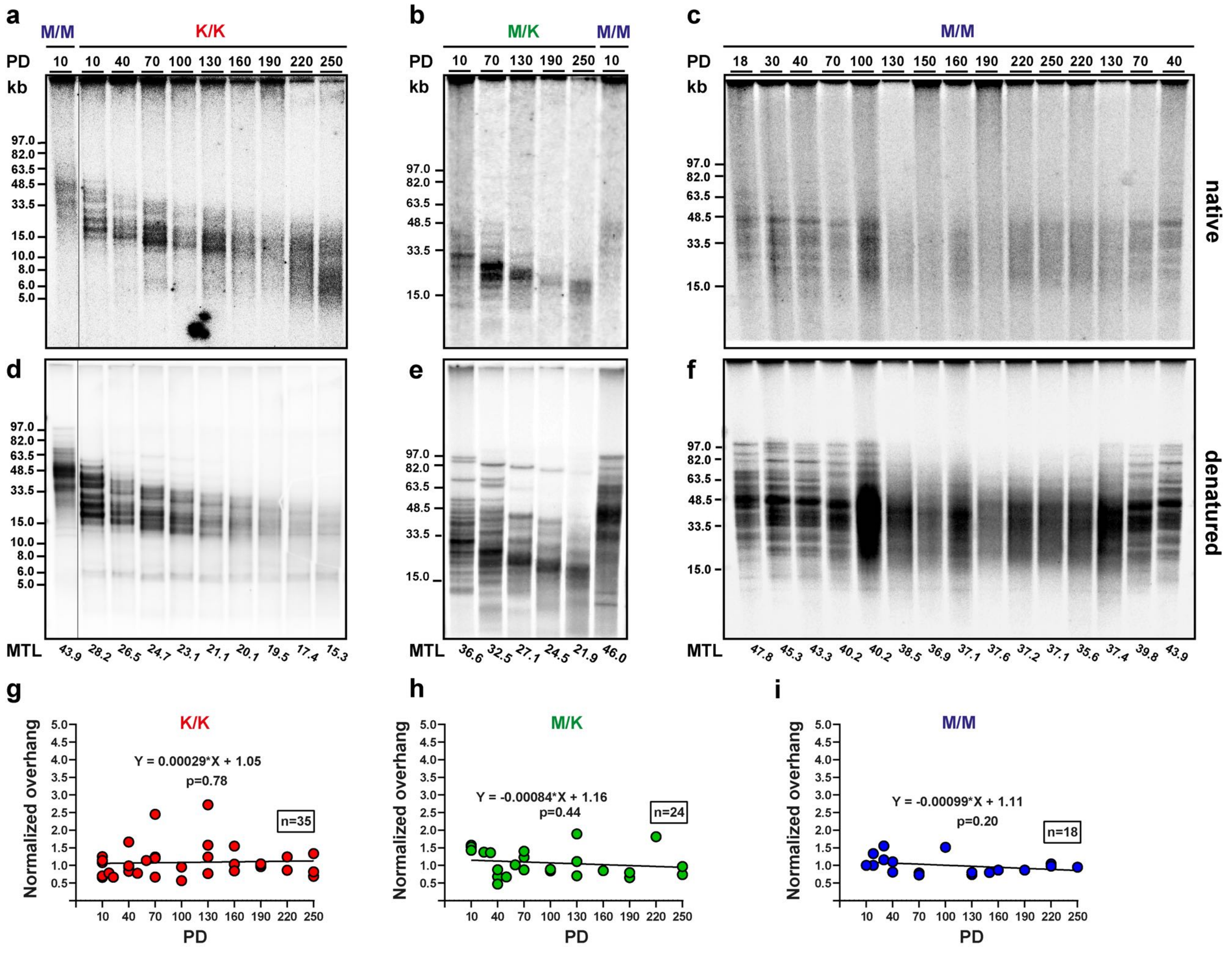


**Figure S7. Normal telomeric overhang signals in *mRtel1* mutant MEFs.** Equal amounts of genomic DNA from the indicated MEF cultures and PD were analyzed by PFGE and in-gel hybridization. (a,b,c) First, a telomeric C-rich probe was hybridized to the native DNA, detecting the single-stranded G-rich telomeric overhang . (d,e,f) Then, the DNA was denatured within the gel and re-hybridized to the same probe to detect the entire telomeric DNA . Representative gels are shown. (g,h,i) The native overhang signal for 77 samples in six gels was measured by *ImageQuant-TL* and plotted (summarized in Table S1). The K/K and M/K samples were normalized to the same WT sample (M/M PD 10) within each gel (g,h). The M/M samples were normalized to WT sample M/M PD 18 within each gel (i). The formula for the regression line and the P value corresponding to the deviation of the slope from 0 are indicated above each graph. The high P values indicate no significant change in the overhang signal over time.

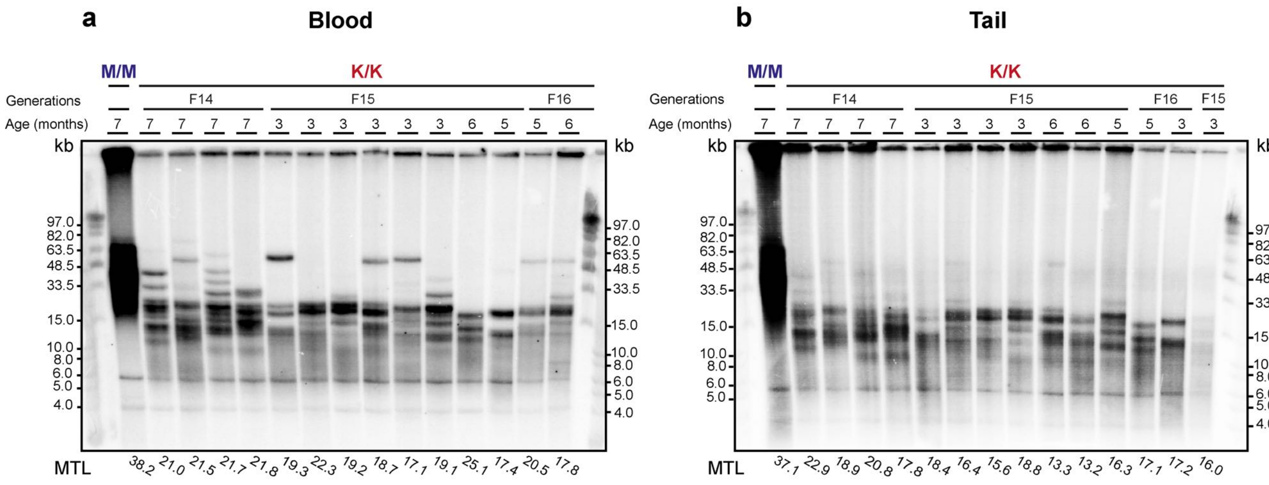


**Figure S8. Telomere length stabilized at late generation Telomice.** Genomic DNA samples extracted from blood leukocytes (a) or tail (b) from 14 Telomice from generations F14, F15 and F16, and one WT (M/M) mouse as a control at the indicated ages, were analyzed by PFGE and in-gel hybridization to the denatured DNA. Mean TRF length for each sample was measured by *Telotool* and indicated below the lanes.

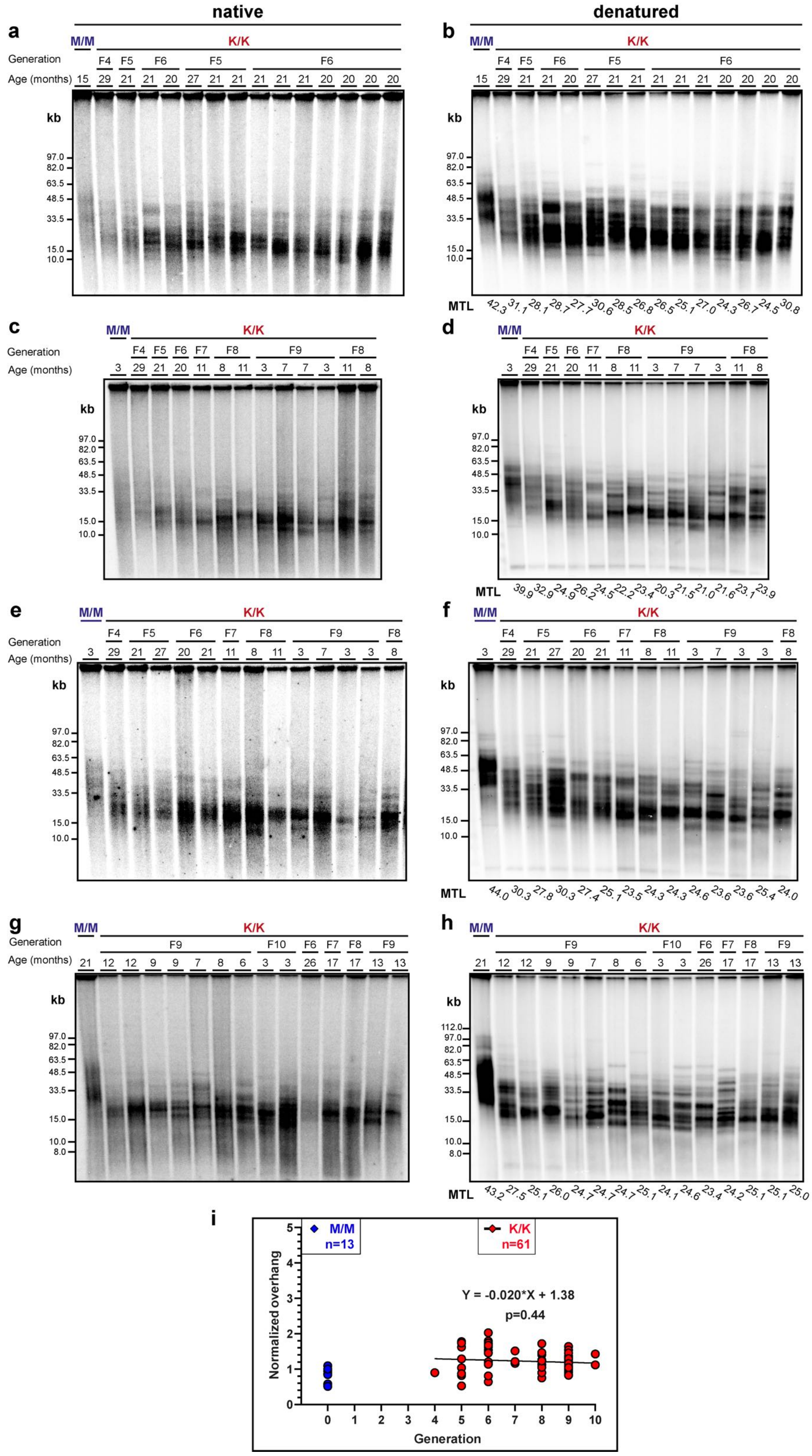

**Figure S9. Normal telomeric overhang signals in tail samples of Telomice.** (a,c,e,g) Equal amounts of genomic DNA derived from tail samples of the indicated Telomice and WT mice were analyzed by PFGE and in-gel hybridizations with a telomeric C-rich probe to the native DNA, detecting the single-stranded G-rich telomeric overhang. (b,d,f,h) The DNA was denatured within the gel and re-hybridized to the same probe to detect the entire telomeric G-strand DNA. Representative gels are shown. (i) The native signal for each sample was measured by *ImageQuant-TL*, and all the samples within each gel (a total of six gels) were normalized to the WT (M/M) sample and plotted (summarized in Table S3). No significant change in the normalized native signal was detected in Telomice tail over seven generations, and no significant difference was found when comparing the Telomice overhang signal to the WT (M/M) mice.

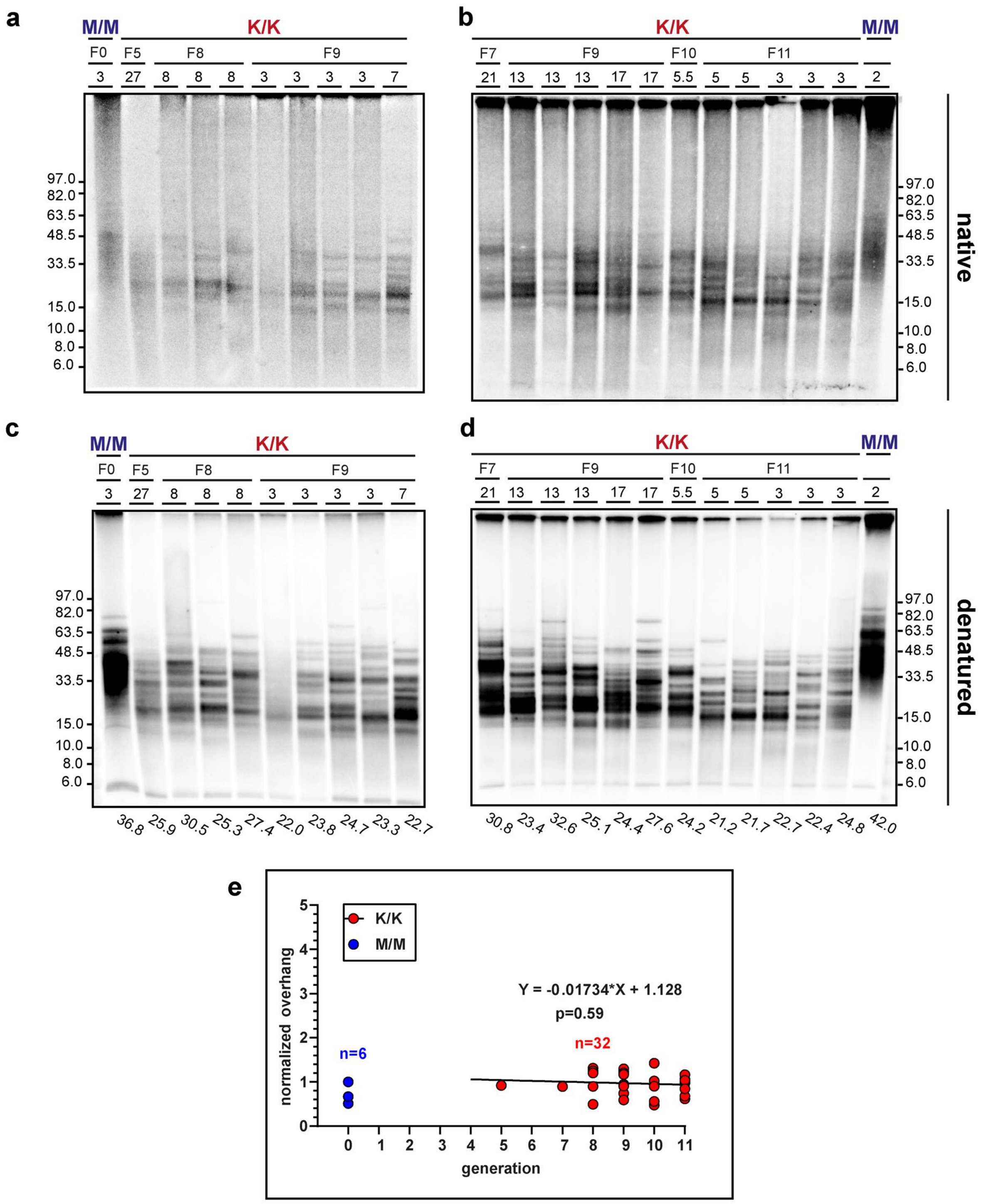


**Figure S10. Normal telomeric overhang signal in blood samples of Telomice.** (a,b) Equal amounts of genomic DNA derived from blood samples of the indicated Telomice and WT mice were analyzed by PFGE and in-gel hybridizations with a telomeric C-rich probe to the native DNA, detecting the single-stranded G-rich telomeric overhang. (c,d) The DNA was denatured within the gel and re-hybridized to the same probe to detect the entire telomeric DNA. Representative gels are shown. (e) The native signal for each sample was measured by *ImageQuant-TL*, and all the samples within each gel (a total of six gels) were normalized to the WT (M/M) sample and plotted (summarized in Table S3). No significant change was detected in the Telomice blood over six generations, and no significant difference was found when comparing the Telomice overhang signal to the M/M (WT) mice.

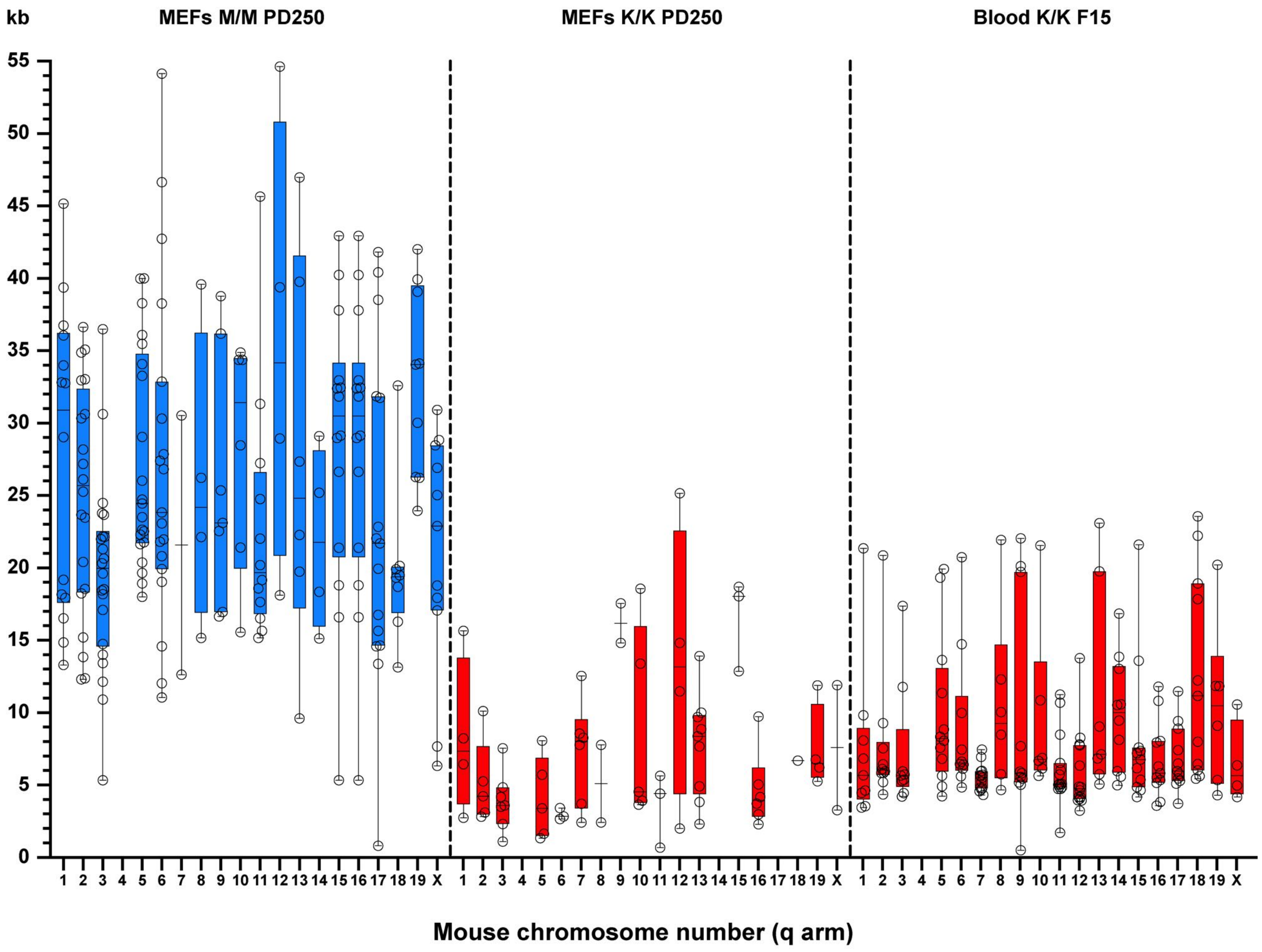


**Figure S11. Mapping of telomere reads to specific mouse chromosome ends.** Individual **t**elomere reads were mapped to the reference mouse genome based on the subtelomeric sequences. Open circles indicate the length of each mapped telomere. Horizontal lines indicate average telomere length, and box boundaries the quartiles.
